# Dopamine projections to the basolateral amygdala drive the encoding of identity-specific reward memories

**DOI:** 10.1101/2022.09.26.509602

**Authors:** Ana C. Sias, Yousif Jafar, Caitlin M. Goodpaster, Kathia Ramírez-Armenta, Tyler M. Wrenn, Nicholas K. Griffin, Keshav Patel, Alexander C. Lamparelli, Melissa J. Sharpe, Kate M. Wassum

**Affiliations:** Dept. of Psychology, UCLA, Los Angeles, CA 90095; Brain Research Institute, UCLA, Los Angeles, CA 90095, USA; Integrative Center for Learning and Memory, University of California Los Angeles, Los Angeles, CA, USA; Integrative Center for Addictive Disorders, University of California Los Angeles, Los Angeles, CA, USA; Department of Psychology, University of Sydney, Camperdown Australia NSW 2006

**Author notes:** Correspondence: Kate Wassum, Dept. of Psychology, UCLA, 1285 Franz Hall, Box 951563, Los Angeles, CA 90095-1563.

**Keywords:** learning, memory, decision making, Pavlovian conditioning, Pavlovian-to-instrumental transfer, ventral tegmental area, basolateral amygdala, appetitive, model-based learning

## Abstract

To make adaptive decisions, we build an internal model of the associative relationships in an environment and use it to make predictions and inferences about specific available outcomes. Detailed, identity-specific cue-reward memories are a core feature of such cognitive maps. Here we used fiber photometry, cell-type and pathway-specific optogenetic manipulation, Pavlovian cue-reward conditioning, and decision-making tests in male and female rats, to reveal that ventral tegmental area dopamine (VTA_DA_) projections to the basolateral amygdala (BLA) drive the encoding of identity-specific cue-reward memories. Dopamine is released in the BLA during cue-reward pairing and VTA_DA_→BLA activity is necessary and sufficient to link the identifying features of a reward to a predictive cue, but does not assign general incentive properties to the cue or mediate reinforcement. These data reveal a dopaminergic pathway for the learning that supports adaptive decision making and help explain how VTA_DA_ neurons achieve their emerging multifaceted role in learning.

---

Dopamine has long been known to critically contribute to learning. Midbrain dopamine neurons can signal errors in reward prediction^1–3^. These learning signals have canonically been interpreted to cache the general value of a reward to its predictor and reinforce response policies that rely on past success, rather than forethought of specific outcomes^1–7^. But adaptive decision making often requires such forethought. For example, if you see both pizza and donut boxes outside the seminar room, assuming you like both, you need to use these cues to represent the identity of the specific predicted foods in order to make the snack choice that is optimal in your current circumstances (e.g., are you craving something sweet or savory?, have you just had donuts for breakfast?). So, to ensure flexible behavior, humans and other animals do not just learn the general value of predictive events, but also encode the relationships between these cues and the identifying features of their associated outcomes^8, 9^. Such identity-specific cue-reward memories are fundamental components of the internal model of environmental relationships, aka cognitive map^10^, we use to generate the predictions and inferences needed for many forms of flexible, advantageous decision making^8, 9, 11, 12^. Little is known of how we form these cue-reward memories. But recent evidence suggests dopamine might actually contribute^13–24^. New data have challenged the value-centric dogma of dopamine function, indicating it plays a much broader role in learning than originally thought^25–29^. How dopamine contributes to identity-specific cue-reward learning is unknown, yet critical for understanding dopamine’s emerging multifaceted function in learning.

One candidate pathway through which dopamine might contribute to cue-reward learning is the VTA dopamine (VTA_DA_) projection to the basolateral amygdala (BLA)^30–35^. This pathway has received much less attention than the more popular VTA_DA_ projections to nucleus accumbens and prefrontal cortex, so little is known of its function. VTA_DA_→BLA projections contribute to Pavlovian fear learning^32^, yet whether this pathway also contributes to appetitive learning and the nature of the memories supported are both unknown. The BLA itself was recently shown to be crucial for forming detailed, identity-specific, cue-reward memories^36^. Therefore, here we combined a systems neuroscience toolkit with Pavlovian cue-reward conditioning and tests of the nature of learning and its influence on decision making to evaluate VTA_DA_→BLA pathway function in linking the unique features of rewarding events to predictive cues, i.e., encoding the identity-specific reward memories that support adaptive decision making.

## RESULTS

### Dopamine is released in the BLA during cue-reward learning

We first asked whether and when dopamine is released in the BLA during the encoding of identity-specific cue-reward memories. We used fiber photometry to record fluorescent activity of the G-protein-coupled receptor-activation-based dopamine sensor GRAB_DA_^37^ in the BLA of male and female rats during Pavlovian conditioning (Figure 1a-c). Rats were food deprived and received 8 sessions of Pavlovian long-delay conditioning during which 2 distinct auditory cues (aka, conditioned stimuli) each predicted a unique food reward (e.g., white noiseꟷsucrose/clickꟷpellets). During each session, each cue was presented 8 times (variable 2.5-min mean intertrial interval, ITI) for 30 s and terminated in the delivery of its associated reward (Figure 1c). This conditioning has been shown to engender the encoding of identity-specific cue-reward memories as evidenced by the ability of the cues to subsequently promote instrumental choice of the specific predicted reward^38–42^ and sensitivity of the conditional goal-approach response to devaluation of the predicted reward^43^. Across training, rats developed a Pavlovian conditional goal-approach response (Figure 1d). Like BLA neuronal responses (Supplemental Figure 1-1), dopamine was released in the BLA at both cue onset and offset/reward delivery across training (Figure 1e-f; see also Supplemental Figure 1-2 for data from each of the 8 training sessions, and Supplemental Figure 1-3 for data aligned to reward collection). Thus, dopamine is released in the BLA in response to both cues and rewarding outcomes, as well as their pairing during Pavlovian conditioning.

**Figure 1.**
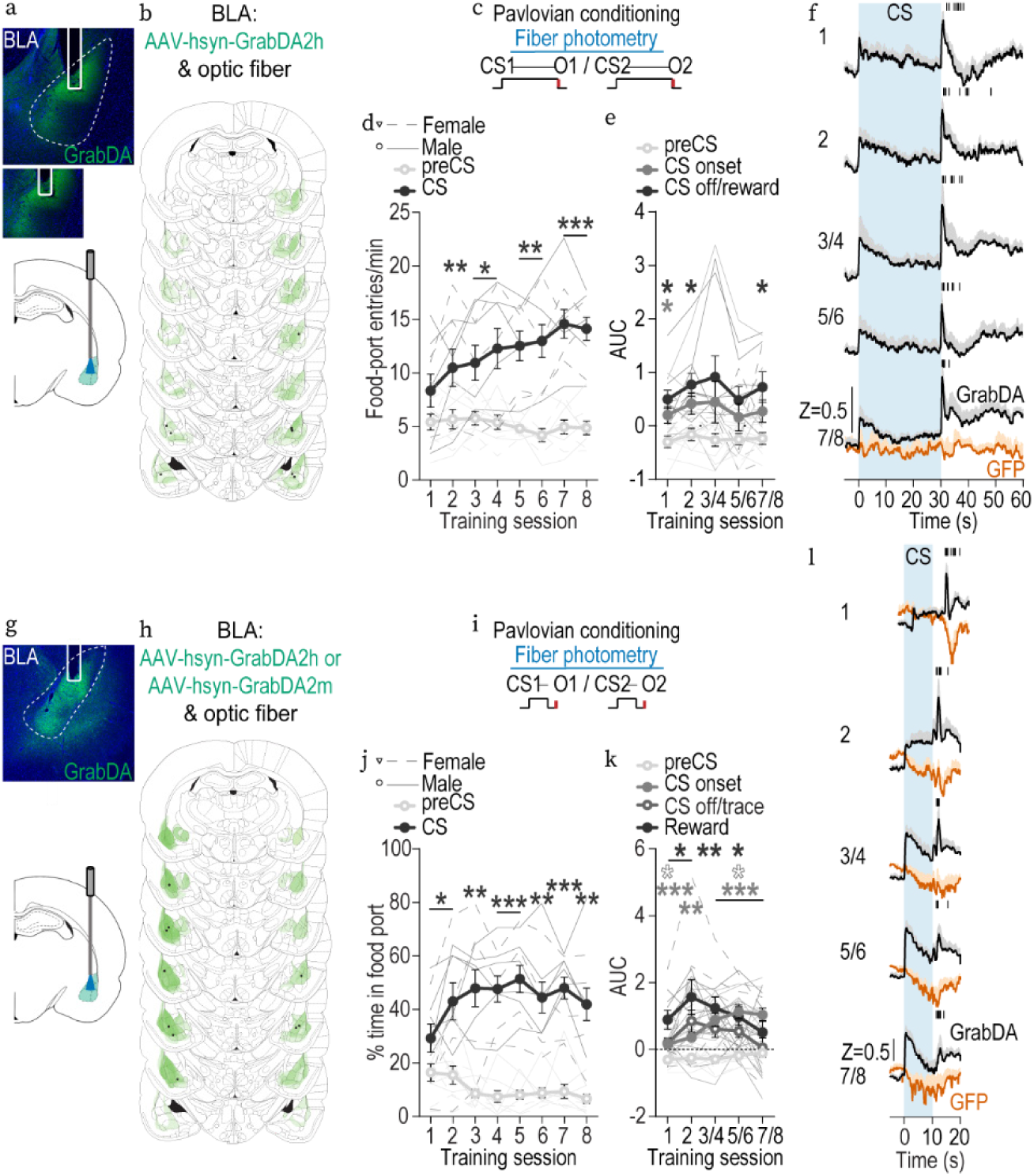
Dopamine is released in the BLA during cue-reward learning. **(a-f)** Fiber photometry recording of dopamine release in the BLA during Pavlovian long-delay conditioning. **(a)** Top: Representative fluorescent image of GRAB_DA2h_ expression and fiber placement in the BLA. Bottom: Schematic of fiber photometry approach for imaging GRAB_DA_ fluorescence changes in BLA neurons. **(b)** Schematic representation of GRAB_DA2h_ expression and placement of optical fiber tips in BLA for all subjects. Brain slides from_115_. **(c)** Pavlovian long-delay conditioning procedure schematic. CS, 30-s conditioned stimulus (aka, “cue”, white noise or click) followed immediately by reward outcome (O, sucrose solution or grain pellet). **(d)** Food-port entry rate during the cue relative to the preCue baseline period, averaged across trials and across the 2 cues for Pavlovian conditioning session. Thin lines represent individual subjects. Training x Cue: *F*_(2.77, 22.15)_ = 14.69, *P <* 0.0001; Training: *F*_(4.75, 38.02)_ = 2.76, *P* = 0.03; Cue: *F*_(1, 8)_ = 44.00, *P =* 0.0002. \**P* < 0.05, \*\**P* < 0.01, \*\*\**P* < 0.001 relative to preCue, Bonferroni correction. **(e)** Trial-averaged quantification of area under the BLA GRAB_DA_ Z-scored curve (AUC) during the 2-s period following CS onset or reward delivery compared to the equivalent baseline period immediately prior to CS onset. Event: F_(1.83, 14.61)_ = 7.63, *P* = 0.006; Training: F_(2.36, 18.85)_ = 1.24, *P* = 0.32; Training x Event: F_(3.72, 29.77)_ = 0.42, *P* = 0.78. * *P* < 0.05, ** *P* < 0.01, relative to preCue baseline, Bonferroni correction. **(f)** Trial-averaged GRAB_DA_ fluorescence changes (Z-score) in response to CS presentation (blue) and outcome delivery across days of training. Shading reflects between-subjects s.e.m. Tick marks represent time of reward collection for each subject. Data from the last six sessions were averaged across 2-session bins (3/4, 5/6, and 7/8). *N* = 9, 5 male. **(g-l)** Fiber photometry recording of BLA dopamine release during Pavlovian trace conditioning. **(g)** Top: Representative fluorescent image of GRAB_DA_ expression and fiber placement in the BLA. Bottom: Schematic of fiber photometry approach for imaging GRAB_DA_ fluorescence changes in BLA neurons. **(h)** Schematic representation of GRAB_DA_ expression and placement of optical fiber tips in BLA for all subjects. **(i)** Pavlovian trace conditioning procedure schematic. CS, 10-s conditioned stimulus (white noise or click) followed by a 1.5-s trace interval before reward outcome (O, chocolate or unflavored purified pellets). **(j)** Percentage of time spent in the food-port during the cue relative to the preCue baseline period, averaged across trials and across the 2 cues for each Pavlovian conditioning session. Thin lines represent individual subjects. Training x Cue: F_(3.21, 28.92)_ = 7.77, *P* = 0.0005; Cue: F_(1, 9)_ = 62.61, *P* < 0.0001; Training: F_(2.80, 25.22)_ = 1.29, *P* = 0.30. \**P* < 0.05, \*\**P* < 0.01, \*\*\**P* < 0.001 relative to preCue, Bonferroni correction. **(k)** Trial-averaged quantification of BLA GRAB_DA_ Z-scored AUC during the 1.5-s period following cue onset, cue offset (trace interval) or reward delivery compared to the equivalent baseline period immediately prior to cue onset. Training x Event: F_(2.85, 25.69)_ = 3.72, *P* = 0.03; Event: F_(1.69, 15.24)_ = 10.80, *P* = 0.002; Training: F_(2.85, 25.63)_ = 2.46, p = 0.09. * *P* < 0.05, ** *P* < 0.01, *** P < 0.001 relative to preCue baseline, Bonferroni correction. **(l)** Trial-averaged GRAB_DA_ fluorescence changes (Z-score) in response to cue presentation and reward delivery across days of training. Shading reflects between-subjects s.e.m. Tick marks represent time of reward collection for each subject. Data from the last six sessions were averaged across 2-session bins (3/4, 5/6, and 7/8). *N* = 10 (GRAB_DA2h_: *N* = 4, 3 male; GRAB_DA2m_: *N* = 6, 3 male).

**Figure 2.**
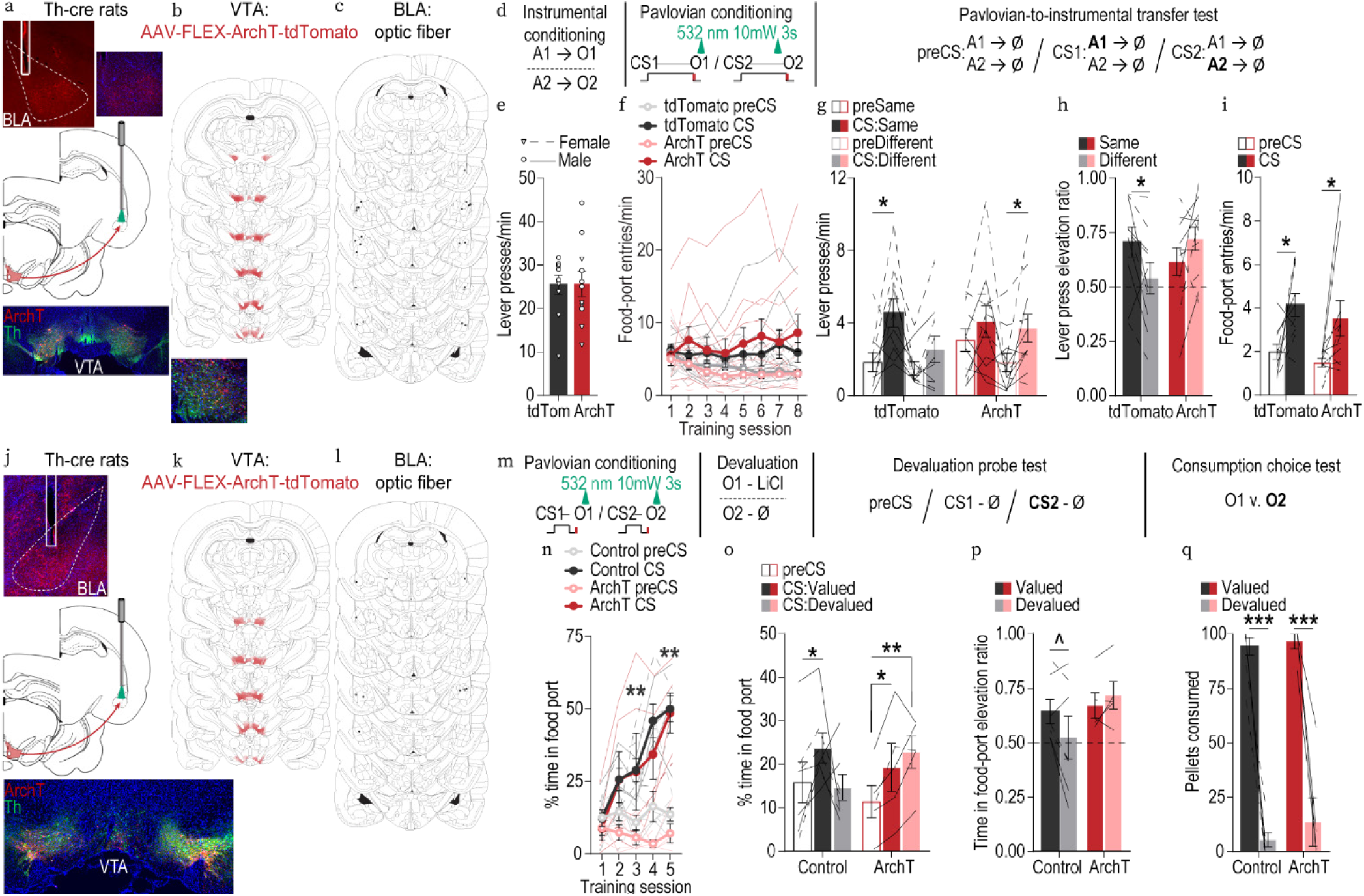
Optical inhibition of VTA_DA_→BLA projections during cue-reward pairing attenuates the encoding of identity-specific cue-reward memories. **(a-i)** Optical inhibition of VTA_DA_→BLA projections during Pavlovian long-delay conditioning with outcome-specific Pavlovian-to-instrumental transfer test. **(a)** Bottom: Representative fluorescent image of cre-dependent ArchT-tdTomato expression in VTA cell bodies with coexpression of Th in Th-Cre rats. Middle: Schematic of optogenetic strategy for bilateral inhibition of VTA_DA_ axons and terminals in the BLA of Th-cre rats. Top: Representative image of fiber placement in the vicinity of immunofluorescent ArchT-tdTomato-expressing VTA_DA_ axons and terminals in the BLA. **(b)** Schematic representation of cre-dependent ArchT-tdTomato expression in VTA and **(c)** placement of optical fiber tips in BLA for all subjects. **(d)** Pavlovian long-delay conditioning and Pavlovian-to-instrumental transfer procedure schematic. A, action (left or right lever press); CS, 30-s conditioned stimulus (aka, “cue”, white noise or click) followed immediately by reward outcome (O, sucrose solution or grain pellet). **(e)** Instrumental conditioning. Lever-press rate averaged across levers and across the final 2 instrumental conditioning sessions. t_(19)_ = 0.07, *P =* 0.94. Data points represent individual subjects. **(f)** Pavlovian conditioning. Food-port entry rate during the cue relative to the preCue baseline period, averaged across trials and across the 2 cues for each Pavlovian conditioning session. Thin lines represent individual subjects. Training x Cue: F_(4.09, 77.71)_ = 5.73, *P =* 0.0004; Cue: *F*_(1,19)_ = 10.34, *P =* 0.005; Training: F_(1.47, 27.98)_ = 1.19, *P =* 0.31; Virus: F_(1,19)_ = 0.05, *P =* 0.83; Training x Virus: F_(7,133)_ = 1.23, *P =* 0.29; Virus x Cue: F_(1,19)_ = 1.04, *P* = 0.32; Training x Virus x Cue: F_(7,133)_ = 0.75, *P* = 0.63. **(g-i)** Outcome-specific Pavlovian-to-instrumental transfer test. **(g)** Trial-averaged lever-press rates during the preCue baseline periods compared to press rates during the cue separated for presses on the lever that, in training, earned the same outcome as predicted by the cue (Same) and pressing on the other available lever (Different). Virus: F_(1, 19)_ = 0.93, *P =* 0.35; Lever: F_(1, 19)_ = 3.36, *P =* 0.08; Cue: F_(1, 19)_ = 22.02, *P =* 0.0002; Virus x Lever: F_(1, 19)_ = 0.12, *P =* 0.73; Virus x Cue: F_(1, 19)_ = 0.37, *P* = 0.55; Lever x Cue: F_(1, 19)_ = 0.25, *P =* 0.62; Virus x Lever x Cue: F_(1, 19)_ = 2.63, *P* =0.12. \**P* < 0.05, planned comparisons CS same presses v. preCue same presses and cue different presses v. preCue different presses. **(h)** Elevation in lever presses on the Same lever [(Same lever presses during cue)/(Same presses during cue + Same presses during preCue)], relative to the elevation in pressing on the Different lever [(Different lever presses during cue)/(Different presses during cue + Different presses during preCue)], averaged across trials and across cues during the PIT test. Virus x Lever: F_(1, 19)_ = 9.22, *P =* 0.007; Virus: F_(1, 19)_ = 0.33, *P =* 0.57; Lever: F_(1, 19)_ = 0.45, *P =* 0.51. \**P* < 0.05, Bonferroni correction. Lines represent individual subjects. **(i)** Food-port entry rate during the cues relative to the preCue baseline, averaged across trials and across the 2 cues during the PIT test. Cue: F_(1, 19)_ = 15.18, *P* = 0.001; Virus: F_(1, 19)_ = 1.15, *P =* 0.30; Virus x Cue: F_(1, 19)_ = 0.008, *P =* 0.93. \**P* < 0.05, Bonferroni correction. ArchT, *N* = 11, 6 males; tdTomato, *N* = 10, 5 males. **(j-q)** Optical inhibition of VTA_DA_→BLA projections during Pavlovian trace conditioning with outcome-specific devaluation test. **(j)** Bottom: Representative fluorescent image of cre-dependent ArchT-tdTomato expression in VTA cell bodies with coexpression of Th in Th-Cre rats. Middle: Schematic of optogenetic strategy for bilateral inhibition of VTA_DA_ axons and terminals in the BLA of Th-cre rats. Top: Representative image of fiber placement in the vicinity of immunofluorescent ArchT-tdTomato-expressing VTA_DA_ axons and terminals in the BLA. **(k)** Schematic representation of cre-dependent ArchT-tdTomato expression in VTA and **(l)** placement of optical fiber tips in BLA for all subjects. **(m)** Pavlovian trace conditioning and outcome-specific devaluation procedure schematic. CS, 10-s conditioned stimulus (white noise or tone) following by 1.5-s trace interval before reward outcome (O, chocolate or unflavored purified pellets); LiCl, lithium chloride 0.3M, 1.5% volume/weight. **(n)** Pavlovian conditioning. Percentage of time spent in the food-delivery port during the cue relative to the preCue baseline period, averaged across trials and across the 2 cues for each Pavlovian conditioning session. Thin lines represent individual subjects. Training x Cue: F_(1.61, 16.13)_ = 31.49, *P* <0.0001; Training: F_(2.82, 28.18)_ = 12.22, *P* < 0.0001; Cue: F_(1, 10)_ = 30.92*,P =* 0.0002; Virus: F_(1, 10)_ = 1.55, *P* = 0.24; Training x Virus: F_(4, 40)_ = 0.87, *P* = 0.49; Virus x Cue: F_(1, 10)_ = 0.23, *P* = 0.64; Training x Virus x Cue: F_(4, 40)_ = 0.30, *P* = 0.88. \*\**P* < 0.01, Bonferroni correction. **(o-p)** Outcome-specific devaluation probe test. **(o)** Trial-averaged percentage of time spent in the food port during the preCue baseline periods compared to that during the Cue signaling the devalued v. non-devalued (valued) reward. Virus x Cue: F_(2, 20)_ = 2.54, *P* = 0.10; Virus: F_(1, 10)_ = 0.003, *P =* 0.95; Cue: F_(1.25, 12.54)_ = 3.14, *P =* 0.09. \**P* < 0.05, \*\**P* < 0.01 planned comparisons preCue v. cue. **(p)** Elevation in percent time spent in food port during the cue signaling the valued reward relative to preCue [(CS Valued % time in port)/(CS Valued % time in port + preCue % time in port)], relative to the elevation in percent time spent in food port during the cue signaling the devalued reward [(CS Devalued % time in port)/(CS Devalued % time in port + preCue % time in port)], averaged across trials and across cues during the devaluation probe test. Virus x Cue: F_(1, 10)_ = 5.20, *P =* 0.046; Virus: F_(1, 10)_ = 1.19, *P =* 0.30; Cue: F_(1, 10)_ = 1.00, *P =* 0.34. ^*P* = 0.059, Bonferroni correction. Lines represent individual subjects. **(q)** Consumption choice test to validate efficacy of devaluation. Amount out of 100 available pellets consumed. Value: F_(1, 10)_ = 249.00, *P* < 0.0001; Virus: F_(1, 10)_ = 0.79, *P =* 0.39; Virus x Value: F_(1, 10)_ = 0.28, *P =* 0.61. \*\*\**P* < 0.001, Bonferroni correction. ArchT, *N* = 5, 4 males; Control, *N* = 7, 4 males (3 WT/cre-dependent ArchT; 4 Th-cre/cre-dependent tdTomato).

**Figure 3.**
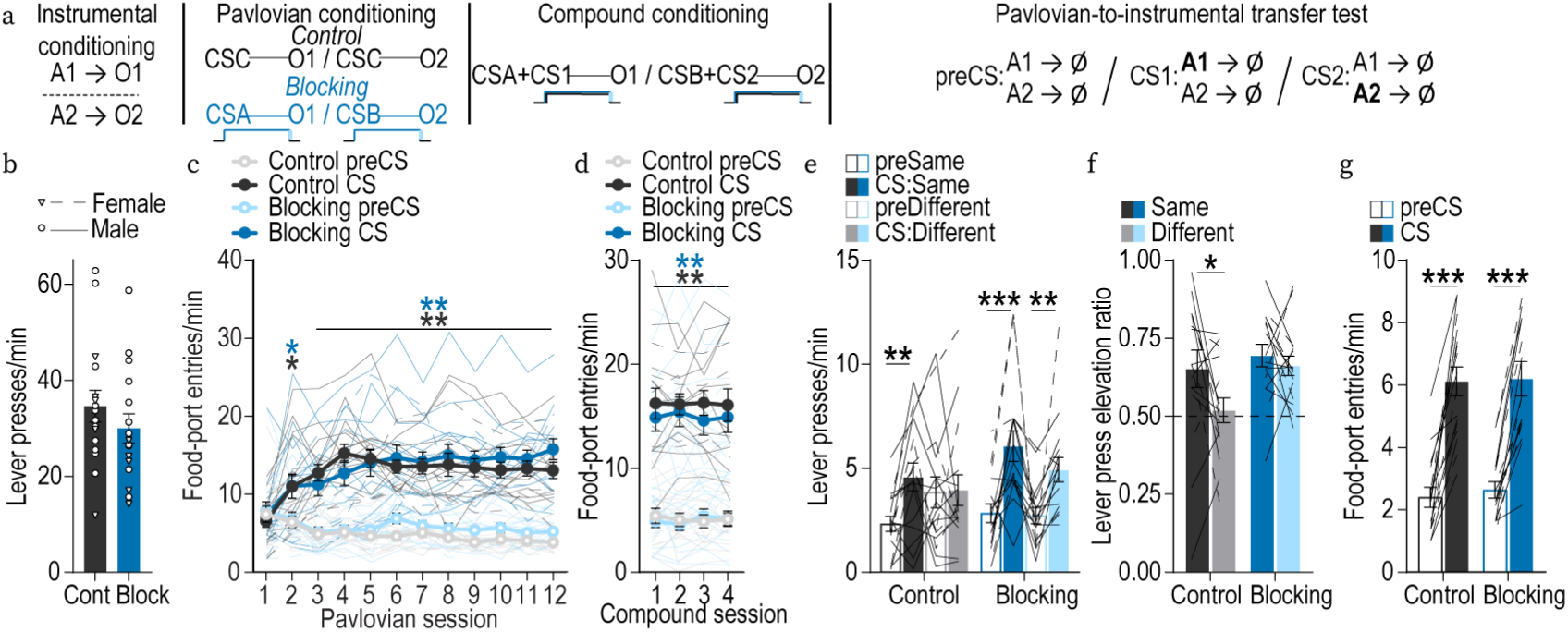
Previously learned cue-reward relationships block encoding of new identity-specific cue-reward memories. **(a)** Procedure schematic. A, action (left or right lever press); CS, 30-s conditioned stimulus (aka, “cue”, CSA/B: house light or flashing lights; CSC: alternating lights on either side of the chamber; CS1/CS2: white noise or click) followed immediately by reward outcome (O, sucrose solution or grain pellet). **(b)** Instrumental conditioning. Lever-press rate averaged across levers and across the final 2 instrumental conditioning sessions. t_(30)_ = 1.03, *P =* 0.31. **(c)** Pavlovian conditioning. Food-port entry rate during the visual cues relative to the preCue baseline periods, averaged across trials and across cues for each Pavlovian conditioning session. Thin lines represent individual subjects. Training x Cue: F_(4.55, 136.40)_ = 30.77, *P* < 0.0001; Training: F_(2.71, 81.41)_ = 4.29, *P =* 0.009; Cue: F_(1, 30)_ = 186.20, *P* < 0.0001; Group: F_(1, 30)_ = 0.56, *P* = 0.46; Training x Group: F_(11, 330)_ = 1.77, p=0.06; Group x Cue: F_(1, 30)_ = 0.22, p=0.64; Training x Group x Cue: F_(11, 330)_ = 0.98, *P* = 0.47. * *P* < 0.05, \*\**P* < 0.01, Bonferroni correction. **(d)** Compound conditioning. Food-port entry rate during the compound cues relative to the preCue baseline, averaged across trials and across the 2 compound cues for each compound conditioning session. Cue: F_(1,30)_ = 173.60, *P* < 0.0001; Training: F_(1.32, 39.71)_ = 0.01, *P =* 0.96; Group: F_(1, 30)_ = 0.35, *P =* 0.56; Training x Group: F_(3, 90)_ = 0.12, *P =* 0.95; Training x Cue : F_(2.50, 75.01)_ = 0.50, *P =* 0.65; Group x Cue: F_(1, 30)_ = 0.51, *P =* 0.48; Training x Group x Cue: F_(3, 90)_ = 0.89, *P* = 0.45. \*\**P* < 0.01, Bonferroni correction **(e-g)** Auditory cue outcome-specific Pavlovian-to-instrumental transfer test. **(e)** Trial-averaged lever-press rates during preCue baseline periods compared to press rates during the auditory cues separated for presses on the lever that, in training, delivered the same outcome as predicted by the auditory cue (Same) and pressing on the other available lever (Different). Group x Cue: F_(1, 30)_ = 4.54, *P =* 0.04; Lever x Cue: F_(1,30)_ = 6.24, *P =* 0.02; Cue: F_(1,30)_ = 29.11, *P* < 0.0001; Group: F_(1, 30)_ = 0.59, *P =* 0.45; Lever: F_(1,30)_ = 0.06, *P =* 0.81; Group x Lever: F_(1, 30)_ = 2.09, *P =* 0.16; Group x Lever x Cue: F_(1, 30)_ = 0.81, *P* = 0.38. \*\**P* < 0.01, \*\*\**P* < 0.001 planned comparisons cue same presses v. preCue same presses and cue different presses v. preCue different presses. **(f)** Elevation in lever presses on the Same lever [(Same lever presses during cue)/(Same presses during cue + Same presses during preCue)], relative to the elevation in presses on the Different lever [(Different lever presses during cue)/(Different presses during cue + Different presses during preCue)], averaged across trials and across cues during the PIT test. Lines represent individual subjects. Group: F_(1, 30)_ = 3.99, *P =* 0.06; Lever: F_(1, 30)_ = 4.35, *P =* 0.046; Group x Lever: F_(1, 30)_ = 1.57, *P =* 0.22. \**P* < 0.05, Bonferroni correction. **(g)** Food-port entry rate during the cues relative to the preCue baseline, averaged across trials and across the 2 cues during the PIT test. Cue: F_(1, 30)_ = 154.70, *P* < 0.0001; Group: F_(1, 30)_ = 0.10, *P =* 0.75; Group x Cue: F_(1, 30)_ = 0.06, *P =* 0.80. \*\*\**P* < 0.001, Bonferroni correction. Blocking, *N* = 16, 11 males; Control, *N* = 16, 11 males.

To further reveal how BLA dopamine relates to cue-reward learning, we recorded GRAB_DA_ in the BLA during Pavlovian trace conditioning (Figure 1g-j). A new group of food-deprived rats received 8 sessions of Pavlovian trace conditioning during which 2 distinct auditory cues each predicted a unique food reward after a brief delay (e.g., white noiseꟷchocolate pellets/clickꟷunflavored pellets; Figure 1i-j). During each session, each cue was presented 8 times (variable 2.5-min ITI) for 10 s and its associated reward was delivered 1.5 s after cue offset. The trace interval temporally separated reward delivery from cue offset, allowing us to resolve dopamine signals to these discrete events. We used a shorter cue-reward interval to better enable subjects to predict reward delivery. This conditioning has also been shown to engender the encoding of identity-specific cue-reward memories as evidenced by sensitivity of the conditional goal-approach response to devaluation of the predicted reward^44^. Again, we found that BLA dopamine is associated with cue-reward learning (Figure 1k-l). BLA dopamine was initially robustly released in response to reward delivery. Thus, across types of cue-reward learning, BLA dopamine is released during cue-reward pairing, the critical window for encoding the cue-reward association. In this task, we found that reward-evoked BLA dopamine attenuated with training. Conversely, cue-evoked BLA dopamine release was initially small and grew with training. Indeed, the slope of the BLA dopamine reward response across training was negative (β = -0.13, confidence interval -0.37 – 0.10) and signifantly different (F_(1,96)_ = 9.09, *P* = 0.003) from the slope of the BLA dopamine cue-onset response across training, which was positive (β = 0.25, confidence interval 0.15 – 0.36). Thus, unpredicted rewards trigger dopamine release in the BLA and this response backpropagates to reward predictors with training. After training, we detected BLA dopamine responses to unpredicted reward delivery, which were graded by reward magnitude (Supplemental Figure 1-4a-b). Consistent with a prior report^34^, we also detected BLA dopamine responses to a mildly aversive event (unpredicted puffs of air to the face; Supplemental Figure 1-4c-d). Thus, dopamine is released in the BLA during salient appetitive and aversive events and during multiple forms of cue-reward learning.

**Figure 4.**
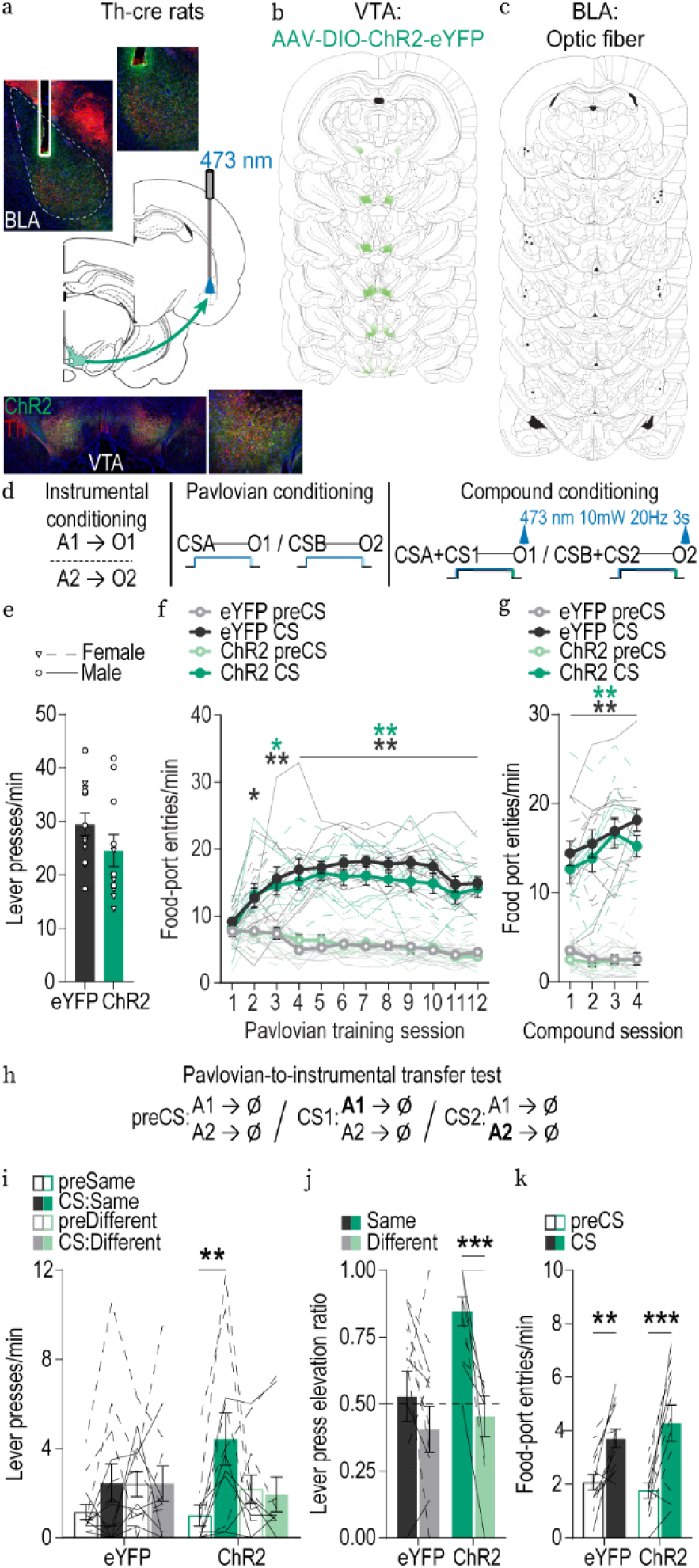
Optical stimulation of VTA_DA_→BLA projections during cue-reward pairing unblocks encoding of identity-specific cue-reward memories. **(a)** Bottom: Representative fluorescent image of cre-dependent ChR2-eYFP expression in VTA cell bodies with co-expression of Th in Th-Cre rats. Middle: Schematic of optogenetic strategy for bilateral stimulation of VTA_DA_ axons and terminals in the BLA. Top: Representative image of fiber placement in the vicinity of immunofluorescent ChR2-eYFP-expressing VTA_DA_ axons and terminals in the BLA. **(b)** Schematic representation of cre-dependent ChR2-eYFP expression in VTA and **(c)** placement of optical fiber tips in BLA for all subjects. **(d)** Schematic of training procedures. A, action (left or right lever press); CS, 30-s conditioned stimulus (aka, “cue”, CSA/B: house light or flashing lights; CS1/CS2: white noise or click) followed immediately by reward outcome (O, sucrose solution or grain pellet). **(e)** Instrumental conditioning. Lever-press rate averaged across levers and across the final 2 instrumental conditioning sessions. t_(22)_ = 1.39, *P =* 0.18. **(f)** Pavlovian conditioning. Food-port entry rate during the visual cues relative to the preCue baseline periods, averaged across trials and across the 2 visual cues for each Pavlovian conditioning session. Thin lines represent individual subjects. Training x Cue: F_(4.15, 91.32)_ = 25.86, *P* < 0.0001; Training: F_(2.60, 57.21)_ = 7.22, *P =* 0.0006; Cue: F_(1, 22)_ = 264.70, *P* < 0.0001; Virus: F_(1, 22)_ = 0.67, *P =* 0.42; Training x Virus: F_(11, 242)_ = 0.47, *P =* 0.92; Virus x Cue: F_(1, 22)_ = 2.24, p=0.15; Training x Virus x Cue: F_(11, 242)_ = 0.86, *P =* 0.58. \**P* < 0.05, \*\**P* < 0.01, Bonferroni correction. **(g)** Compound conditioning. Food-port entry rate during the compound cues relative to the preCue baseline, averaged across trials and across the 2 compound cues for each compound conditioning session. Training x Cue period: F_(2.28, 50.21)_ = 9.06, *P =* 0.0002; Training: F_(1.35, 29.67)_ = 6.43, *P =* 0.01; Cue : F_(1, 22)_ = 232.10, *P* < 0.0001; Virus: F_(1, 22)_ = 0.88, *P =* 0.36; Training x Virus: F_(3, 66)_ = 1.01, *P =* 0.40; Virus x Cue: F_(1, 22)_ = 0.54, *P =* 0.47; Training x Virus x Cue: F_(3, 66)_ = 1.07, *P* = 0.37 \*\**P* < 0.01, Bonferroni correction. **(h-k)** Auditory cue outcome-specific Pavlovian-to-instrumental transfer test. **(h)** Test procedure schematic. **(i)** Trial-averaged lever-press rates during the preCue baseline compared to press rates during the auditory cues separated for presses on the lever that, in training, delivered the same outcome as predicted by the auditory cue (Same) and pressing on the other available lever (Different). Virus x Lever x Cue: F_(1, 22)_ = 4.48, *P =* 0.046; Lever x Cue: F_(1, 22)_ = 19.04, *P =* 0.0002; Lever: F_(1, 22)_ = 0.001, *P =* 0.97; Virus: F_(1, 22)_ = 0.14, *P =* 0.72; Cue: F_(1, 22)_ = 7.45, *P =* 0.01; Virus x Lever: F_(1, 22)_ = 1.57, p =0.22; Virus x Cue: F_(1, 22)_ = 1.24, *P =* 0.28. \*\**P* < 0.01, planned comparisons cue same presses v. preCue same presses and cue different presses v. preCue different presses. **(j)** Elevation in presses on the Same lever [(Same lever presses during cue)/(Same presses during cue + Same presses during preCue)], relative to the elevation in responding on the Different lever [(Different lever presses during cue)/(Different presses during cue + Different presses during preCue)], averaged across trials and across cues during the PIT test. Lines represent individual subjects. Virus x Lever: F_(1, 22)_ = 5.72, *P =* 0.03; Virus: F_(1, 22)_ = 3.29, *P =* 0.08; Lever: F_(1, 22)_ = 20.82, *P =* 0.0002. \*\*\**P* < 0.001, Bonferroni correction. **(k)** Food-port entry rate during the cue relative to the preCue period, averaged across trials and across the 2 cues during the PIT test. Cue: F_(1, 22)_ = 36.10, *P* < 0.0001; Virus: F_(1, 22)_ = 0.08, *P =* 0.77; Virus x Cue: F_(1, 22)_ = 1.65, *P =* 0.21. \*\**P* < 0.01, \*\*\**P* < 0.001, Bonferroni correction. ChR2, *N* = 11, 6 males; eYFP, *N* = 13, 6 males.

### VTA_DA_→BLA projections are necessary for encoding identity-specific cue-reward memories

Having found that dopamine is released in the BLA during cue-reward pairing, we next asked whether this mediates the encoding of identity-specific cue-reward memories (Figure 2a-d). We cre-dependently expressed the inhibitory opsin archaerhodopsin T (ArchT) or tdTomato control bilaterally in VTA_DA_ neurons of male and female tyrosine hydroxylase (Th)-cre rats^45^ (Figure 2a-b) and implanted optical fibers bilaterally over BLA (Figure 2c) to allow us to, in ArchT-expressing subjects, transiently inactivate VTA_DA_ axons and terminals in the BLA. Rats first received instrumental conditioning, without manipulation, in which one of two different lever-press actions each earned one of two distinct food rewards (e.g., left press→sucrose/right press→pellets; 11 sessions; Figure 2e). Rats then received Pavlovian long-delay conditioning, during which each of 2 distinct, 30-s, auditory cues predicted the immediate delivery of one of the food rewards at cue offset (e.g., white noiseꟷsucrose/clickꟷpellets; 8 of each cue/session; variable 2.5-min mean ITI; 8 sessions). VTA_DA_→BLA projections were optically inhibited (532 nm, 10 mW, 3 s) coincident with each reward delivery during each Pavlovian conditioning session. We restricted optical inhibition to reward delivery because this is the time at which the cue-reward pairing occurs and when we detected robust dopamine release in the BLA (Figure 1f). Optical inhibition of VTA_DA_→BLA projections did not disrupt outcome collection (Supplemental Figure 2-1a). It also did not impede the development of a Pavlovian conditional goal-approach response (Figure 2f), even if we inhibited throughout the entire cue and reward period (Supplemental Figure 2-2). Thus, VTA_DA_→BLA projections are not required to reinforce an appetitive conditional goal-approach response.

Conditional approach to the shared goal location does not require subjects to have learned the identifying details of the predicted rewards. So, to ask whether VTA_DA_→BLA projections are needed for encoding identity-specific cue-reward memories, we next gave subjects an outcome-specific Pavlovian-to-instrumental transfer (PIT) test. During this test both levers were present, but lever pressing was not reinforced. Each cue was presented 4 times (also without accompanying reward), with intervening cue-free baseline periods (fixed 2.5-min ITI), to assess its influence on action performance and selection in the novel choice scenario. Because the cues are never directly associated with the instrumental actions, this test assesses the ability to use the cues to retrieve a representation of the specific predicted reward to motivate choice of the action known to earn that same unique outcome^39, 46, 47^. No manipulation was given on test. If subjects had encoded identity-specific cue-reward memories, then cue presentation should cause them to selectively increase presses on the lever that, during training, earned the same specific reward as predicted by that cue. Controls showed this outcome-specific PIT effect, increasing presses during the cues on the lever associated with the same predicted reward, but not on the lever associated with the different reward. Conversely, the cues were not capable of selectively guiding lever-press choice in the group for which VTA_DA_→BLA projections were inhibited during Pavlovian conditioning (Figure 2g-h). Rather, for these subjects, the cues increased pressing on both levers, significantly so on the different lever, suggesting the general incentive properties of the cues were intact. Indeed, inhibition of VTA_DA_→BLA projections during the entire cue and reward period during learning, prevented encoding of identity-specific cue-reward memories, but still did not disrupt the ability of cues to non-discriminately motivate instrumental action (Supplemental Figure 2-2). As in training, during the PIT test the conditional goal-approach response was similar between groups (Figure 2I; see also Supplemental Figure 2-2 for similar results with longer duration inhibition). Thus, VTA_DA_→BLA projections are active at the time of cue-reward pairing and this activity is needed to link the identifying details of the reward to the predictive cue, but not to reinforce a conditional response or to assign general incentive properties to the cue to support general motivation.

To provide converging evidence that VTA_DA_→BLA projections support the encoding of identity-specific cue-reward memories, we next asked whether this pathway mediates the cue-reward learning that enables subjects to adapt their conditional responses following devaluation of the predicted reward (Figure 2a-m). We used the Pavlovian trace conditioning task to further generalize VTA_DA_→BLA function across different types of cue-reward learning. During each conditioning session, each of 2 distinct, 10-s, auditory cues was presented 8 times for 10 s and its associated reward was delivered 1.5 s after cue offset (e.g., white noiseꟷchocolate pellets/pulsed toneꟷunflavored pellets; 5 sessions; Figure 2m). VTA_DA_→BLA projections were optically inhibited (532 nm, 10 mW, 3 s) with each reward delivery to attenuate the BLA dopamine reward response (Figure 1l) during learning. Again, optical inhibition of VTA_DA_→BLA projections did not disrupt reward collection (Supplemental Figure 2-1b) or impede the development of a Pavlovian conditional goal-approach response (Figure 2n), confirming VTA_DA_→BLA projections are not required to reinforce an appetitive Pavlovian response.

To ask whether VTA_DA_→BLA are needed to encode identity-specific cue-reward memories, we next devalued one of the food rewards by pairing it with lithium chloride (LiCl; 0.3M, 1.5% volume/weight; 8 sessions) in the absence of the associated cue. The devaluation was effective. Rats fully rejected the LiCl-paired but not unpaired food (Figure 2q). During test, each cue was presented 8 times (without accompanying reward), with intervening cue-free baseline periods (variable 2.5-min ITI). No manipulation was given on test. If subjects had encoded identity-specific cue-reward memories, they will use the cues to retrieve a representation of the specific predicted reward and increase entries into the food-port during the cue signaling the non-devalued reward but not during the cue signaling the devalued reward^9, 48, 49^. Controls showed this outcome-specific devaluation effect. Conversely, the conditional food-port approach response of subjects for which we inhibited VTA_DA_→BLA projections learning was insensitive to devaluation (Figure 2o-p). Rats continued to spend time in food port during the cue, even if the predicted outcome was devalued. Combined these data indicate that dopamine is released in the BLA during reward experience and this is necessary to link the identifying features of the reward to a predictive cue to enable subsequent cue-induced reward predictions for adaptive decision making.

### VTA_DA_**→**BLA projection activity is sufficient to drive the encoding of identity-specific cue-reward memories

Since VTA_DA_→BLA projection activity during cue-reward pairing mediates the encoding of identity-specific cue-reward memories, we reasoned that activation of these projections might drive the formation of such memories. To test this, we first needed to attenuate the encoding of cue-reward memories to serve as a platform to neurobiologically rescue learning. To achieve this, we took advantage of classic Kamin blocking procedures^50–52^ by using a visual cue that already reliably predicts a particular reward to block formation of an association between a novel auditory cue and that specific reward^53^ (Figure 3a). Male and female rats first received instrumental conditioning in which each of two lever-press actions earned a unique food reward (e.g., left press→sucrose/right press→pellets; 11 sessions; Figure 3b). Subjects then received visual cue Pavlovian conditioning. For subjects in the Blocking group, two distinct 30-s visual cues each terminated in the delivery of a unique food outcome (e.g., house lightꟷsucrose/flashing lightꟷpellet; 16 of each cue/session; 2.5-min mean variable ITI; 12 sessions). Controls received equated conditioning in which a third distinct 30-s visual stimulus predicted both reward types (16 trials alternating lights-sucrose/16 trials alternating lights-pellet). Subjects acquired Pavlovian conditional goal-approach responses to these visual cues (Figure 3c). All subjects then received compound conditioning, during which each of the two visual cues previously conditioned for the Blocking group was presented concurrent with an auditory cue for 30 s terminating in the delivery of one of the distinct food outcomes (e.g., house light + white noiseꟷsucrose/flashing light + clickerꟷpellet; 8 of each compound cue/session; 4 sessions). For subjects in the blocking group, each compound cue was paired with the reward previously associated with the visual cue. Thus, the visual component of the compound cue already reliably predicted the outcome. However, for controls neither the visual nor auditory component of the compound cue had been previously associated with the outcome. All subjects showed conditional goal-approach responses to the compound cues (Figure 3d). To assess acquisition of the unique, identity-specific auditory cue-reward relationships, rats were given a PIT test during which action selection was evaluated in the presence of the auditory cues. Controls showed evidence that they learned the auditory cue-reward relationships by being able to use the cues to selectively increase presses on the lever associated with the same specific reward (Figure 3e-f). If the previously encoded visual cue-reward memory blocked encoding of the relationship between the auditory cue and identifying features of the reward, then subjects in the blocking group should not be able to use the auditory cues to represent the specific predicted reward and guide their choices towards the action associated with that outcome during the PIT test^53^. This is what we found. Subjects in the blocking group displayed a non-specific increase in pressing across both levers during the PIT test, indicating they were unable to use the auditory cues to represent the specific predicted reward to guide choice and, thus, had not learned the identity-specific auditory cue-reward memories (Figure 3e-f). Despite disrupted PIT performance, expression of conditional goal-approach response was preserved in the blocking group (Figure 3g). Thus, as has been shown previously^53^, we were able to effectively attenuate the encoding of identity-specific cue-reward memories.

Using this blocking procedure, we next asked whether activation of VTA_DA_→BLA projections is sufficient to rescue, or unblock, the encoding of identity-specific cue-reward memories (Figure 4a-d). We cre-dependently expressed the excitatory opsin channelrhodopsin (ChR2) or eYFP control in VTA_DA_ neurons of male and female Th-cre rats (Figure 4a-b) and implanted optical fibers bilaterally over BLA (Figure 4c) to allow us to, in ChR2-expressing subjects, transiently stimulate VTA_DA_ axons and terminals in the BLA. Rats first received instrumental conditioning, without manipulation, to learn two action-reward relationships (e.g., left press→sucrose/right press→pellets; Figure 4e). They then received visual cue Pavlovian conditioning, also manipulation-free. All subjects received blocking conditions and, thus, during Pavlovian conditioning had two distinct visual cues each paired with a unique food outcome (e.g., house lightꟷsucrose/flashing lightꟷpellet). Both groups developed Pavlovian conditional goal-approach responses to the visual cues (Figure 4f). Rats next received compound conditioning during which each of the visual cues was presented concurrent with an auditory cue for 30 s terminating in the delivery of the same outcome already associated with the visual cue (e.g., house light + white noiseꟷsucrose/flashing light + clickerꟷpellet). During each compound conditioning session, VTA_DA_→BLA projections were optically stimulated (473 nm; 20 Hz, 10 mW, 25-ms pulse width, 3 s) during reward delivery, when the cue-reward pairing happens and, thus, learning can occur. VTA_DA_→BLA stimulation had no effect on reward collection (Supplemental Figure 4-1). It also did not affect goal-approach responses to the compound cue (Figure 4g). To ask about the encoding of identity-specific cue-reward memories, we gave rats a PIT test with the auditory cues, without manipulation. We replicated the blocking in the eYFP controls. For these subjects, the auditory cues were not capable of guiding choice behavior during the PIT test. Stimulation of VTA_DA_→BLA projections during compound training did, however, drive the encoding of identity-specific cue-reward memories. Rats in this group were able to use the auditory cues to know which specific outcome was predicted to selectively increase presses on the lever associated with that same reward (Figure 4h-i). Both groups showed similar goal-approach responses to the cues (Figure 4j), indicating that optical stimulation of VTA_DA_→BLA projections did not augment reinforcement of a general conditional approach response. Similarly, rats did not self-stimulate VTA_DA_→BLA projections, indicating stimulation at this frequency, which reflects the upper endogenous firing rate of dopamine neurons in response to rewarding events^2, 3, 54^, was not itself reinforcing (Supplemental Figure 4-2). Thus, activation of VTA_DA_→BLA projections concurrent with reward experience is sufficient to drive the encoding of identity-specific cue-reward memories, but does not promote reinforcement.

## DISCUSSION

Here we explored the function of dopamine input to the BLA in cue-reward learning. We found that dopamine is released in the BLA during cue-reward pairing and can backpropagate from reward to predictors with learning. VTA_DA_→BLA projection activity at cue-reward pairing is both necessary and sufficient to drive the encoding of identity-specific, cue-reward memories to enable subsequent adaptive decision making. It does not, however, mediate reinforcement or assign general incentive properties to cues to support non-specific motivation. These data reveal the VTA_DA_→BLA pathway as a critical contributor to the formation of detailed, identity-specific, cue-reward memories, fundamental components of the internal model of environmental relationships, aka cognitive map, that supports flexible decision making.

Dopamine is released in the BLA during cue-reward learning. Across two different forms of cue-reward pairing, we detected robust dopamine responses to reward delivery. Thus, dopamine is released in the BLA when a cue-reward association can be formed and when the BLA is, itself, active^36, 55–60^. During long-delay conditioning, cue-reward pairing continued to be accompanied by dopamine release throughout training, perhaps owing to the cooccurrence of cue offset and reward delivery and/or the difficulty precisely timing reward delivery with a long cue-reward interval. With a shorter cue-reward interval and temporal separation of reward from cue offset, we found that the reward-evoked BLA dopamine response attenuated with learning. Correspondingly, it backpropagated to the reward predictor with training. VTA_DA_ neurons are well known to support learning by signaling errors in reward prediction^1–3^. The backpropagating pattern is consistent with such a signal, as is the finding that dopamine responses to unpredicted reward scaled with reward magnitude. However, a pure reward prediction error signal would dip in response to aversive events^61^. To the contrary, but consistent with the activity of VTA_DA_→BLA terminals^34^, we found that an unpredicted aversive event increased BLA dopamine release. Thus, dopamine is released in the BLA in response to both unpredicted rewarding and aversive events and, therefore, at least in bulk, does not signal valence or solely reward prediction error. We also detected small dopamine responses to the cues when they were novel on the first conditioning session. Combined these findings are consistent with a model in which dopamine integrates salience and reward prediction error^62^ and with evidence that dopamine can reflect perceived salience to support the attention needed for learning^63, 64^. Indeed, BLA dopamine can shape attention-related learning signals in the BLA^65^. VTA_DA_ neurons have recently been implicated in myriad learning-related processes^16, 25–29, 63, 64, 66^. Thus, further work is needed to reveal the precise processes that BLA dopamine encodes to support learning, including the possibility that BLA dopamine release may be heterogeneous based on individual VTA_DA_→BLA cell activity, microenvironment, and/or type of learning. Critically, dopamine is released in the BLA to cues, rewards, and their pairing, when cues can become linked to the identifying features of the rewards they predict.

VTA_DA_→BLA pathway activity at the time of cue-reward pairing drives the encoding of identity-specific reward memories. Inhibiting this activity attenuated the ability to link the identifying details of the reward to the predictive cue such that subjects were unable to later use information to inform decision making in a novel situation. This was supported across two different forms of cue-reward learning (long-delay and trace conditioning) and two different types of decision making (instrumental choice and sensitivity to outcome devaluation). Stimulation of VTA_DA_→BLA projections was sufficient to rescue the encoding of identity-specific cue-reward memories when it would otherwise be blocked, such that subjects were later able to use these memories to inform decision making. Although the terminal inhibition indicates the VTA_DA_→BLA pathway regulates identity-specific reward learning, if VTA_DA_→BLA neurons collateralize, the stimulation results could, in part, be due to antidromic activation of collaterals triggering dopamine release elsewhere in the brain. Nonetheless, the data indicate that VTA_DA_→BLA activity is both necessary and sufficient for the formation of identity-specific reward memories. This is consistent with prior evidence that VTA_DA_ neurons track learning from unexpected changes in outcome identity^67^ and can signal the identifying features of an reward^18^. It also accords with evidence that VTA_DA_ neuron activity mediates unblocking driven by changes in outcome identity^22^ and drives learning about the identifying features of predicted rewards needed for sensitivity of cue responses to outcome devaluation^21^. The present data indicate VTA_DA_ neurons mediate the encoding of identity-specific reward memories and that this is achieved, at least in part, via projections to BLA.

The VTA_DA_→BLA pathway does not mediate reinforcement or assign general incentive properties to cues. The canonical theory of dopamine function is that it provides a teaching signal to cache the general value of future rewarding events to a predictive cue and reinforce response policies based on past success^1–7^. If VTA_DA_→BLA projections mediate reinforcement, then we should have found their inhibition to disrupt the development of the Pavlovian conditional approach response. To the contrary, conditional responses were preserved following VTA_DA_→BLA inhibition. If VTA_DA_→BLA pathway activity is sufficient to promote reinforcement, then we should have found activation of this pathway to be reinforcing itself or to promote the reinforcement of conditional responses. We found no evidence of this either. Rather, VTA_DA_→BLA projections regulate the cue-reward learning that enables subsequent choices in new situations, absent any prior opportunity for those choices to have been reinforced. Two pieces of evidence indicate that VTA_DA_→BLA projection activity does not cache general incentive properties to cues. First, following inhibition of VTA_DA_→BLA projections during learning, cues were still capable of generally invigorating instrumental activity, akin to general Pavlovian-to-instrumental transfer^47, 68–71^. Second, stimulating VTA_DA_→BLA projections during cue-reward pairing did not cause cues to non-discriminately motivate action. These null effects on reinforcement and general motivational value are consistent with evidence that the BLA itself is dispensable for these processes^36, 72, 73^. Thus, any contribution of dopamine to general value and reinforcement learning is likely via pathways other than those to the BLA. VTA_DA_→BLA projections may be specialized for encoding the identity-specific memories that support adaptive decision making.

By establishing a function for the VTA_DA_→BLA pathway in identity-specific reward memory, these data open new and important questions for future investigation. One is the mechanism through which VTA_DA_→BLA projections contribute to learning. Dopamine is positioned to influence learning via modulation of neuronal plasticity in the BLA. Dopamine can act on GABAergic interneurons to increase spontaneous inhibitory network activity^34, 74, 75^ and enhance long-term potentiation through suppression of feedforward inhibition^76^. Like dopamine function in the prefrontal cortex^77, 78^, this balance could enhance signal-to-noise by filtering out weak inputs to ensure only strong inputs conveying important information are potentiated. Dopamine can also enhance the excitability of BLA projection neurons^74^ and activation of VTA_DA_→BLA projections can elevate the second messenger cyclic adenosine monophosphate and enhance BLA responses to cues^79^. Dopamine may gate plasticity in BLA^79–83^, as it does in striatal circuits^84–86^. Separate populations of BLA neurons can encode unique rewards^87, 88^. VTA_DA_→BLA projections may contribute to identity-specific associative learning by facilitating the formation of these neuronal groups. This is a ripe question for future investigation. Another is the excitatory synapses that dopamine signaling may potentiate. One candidate is lateral orbitofrontal cortex projections to the BLA, which mediate the encoding of identity-specific reward memories^36^. At least in mice, some VTA_DA_→BLA projections can corelease glutamate to activate BLA interneurons^34^. That BLA dopamine release coincides with cue-reward pairing suggests dopamine is likely to be involved, but the extent to which glutamate corelease contributes is another important open question.

Findings from this study have important implications for how we conceptualize VTA_DA_ function. We found that dopamine release in the BLA contributes to the learning that enables flexible decision making in novel situations. These results contribute to the emerging understanding that VTA_DA_ neurons have a multifaceted role in learning^25–29^. This and other recent work on more canonical dopamine pathways^21, 89,90^ indicates dopamine’s multifaceted contribution to learning is likely dictated by the function of downstream target regions. As we further explore the function of distinct dopamine pathways we may reveal core principles of dopamine function, e.g., learning and/or plasticity modulation, but we will most certainly find diversity of function based on projection target.

Here we show that the VTA_DA_→BLA pathway drives the formation of an association between a cue and the unique reward it predicts. Such identity-specific reward memories are fundamental components of the internal model of environmental relationships, cognitive map, that enables us to generate the predictions and inferences needed for flexible decision making, including that in novel situations^8, 9, 11, 12^. This core form of memory can support a diverse array of behavioral and decision functions. Thus, VTA_DA_→BLA projections may also support identity-specific social, drug, and/or aversive memories. An inability to properly encode predicted outcomes can lead to ill-informed motivations and decisions. This is characteristic of the cognitive symptoms underlying many psychiatric diseases^91–102^. Thus, these data may also aid our understanding and treatment of substance use disorder and mental illnesses marked by disruptions to both dopamine function and decision making.

## ACKNOWLEDGEMENTS

This research was supported by NIH grant DA035443 and MH106972 (KMW), NIH grant DA057084 (KMW & MJS), NSF GRFP (ACS), NSF CAREER 2143910 (MJS), and the Staglin Center for Behavior and Brain Sciences.

## AUTHOR CONTRIBUTIONS

KMW and ACS designed the research, analyzed, and interpreted the data. ACS conducted the research with assistance from YXJ, NKG, and ACL. CMG and TMW conducted the behavioral blocking experiments. KR contributed to the fiber photometry experiments. NKG and KP assisted with histological verification. MJS contributed to the design of the blocking experiments and advised on the project and manuscript. ACS and KMW wrote the manuscript.

## COMPETING FINANCIAL INTERESTS

The authors declare no biomedical financial interests or potential conflicts of interest.

## METHODS

### Subjects

Male and female wildtype Long-Evans rats and transgenic Long-Evans rats expressing Cre recombinase under control of the tyrosine hydroxylase promoter (Th-cre) aged 8 - 11 weeks at the time of surgery served as subjects. Rats were housed in a temperature (68-79°F) and humidity (30-70%) regulated vivarium. They were initially housed in same-sex pairs and then following surgery housed individually to preserve implants. Rats were provided with water *ad libitum* in the home cage and were maintained on a food-restricted 12-14 g daily diet (Lab Diet, St. Louis, MO) to maintain approximately 85-90% free-feeding body weight. Rats were handled for 3-5 days prior to the onset of each experiment. Separate groups of naïve rats were used for each experiment. Experiments were performed during the dark phase of a 12:12 hr reverse dark/light cycle (lights off at 7AM). All procedures were conducted in accordance with the NIH Guide for the Care and Use of Laboratory Animals and were approved by the UCLA Institutional Animal Care and Use Committee.

### Surgery

We used standard surgical procedures described previously^36, 103–105^. Rats were anesthetized with isoflurane (4–5% induction, 1–2% maintenance), and a nonsteroidal anti-inflammatory agent was administered pre- and postoperatively to minimize pain and discomfort. Surgical details for each experiment are described below. In all cases, surgery occurred prior to the onset of behavioral training.

### Behavioral procedures

#### Apparatus

Training took place in Med Associates conditioning chambers (East Fairfield, VT) housed within sound- and light-attenuating boxes, described previously^106^. Each chamber had grid floors and contained 2 retractable levers that could be inserted to the left and right of a recessed food-delivery port (magazine) on the front wall. Stimulus lights were positioned above each of these levers. A photobeam entry detector was positioned at the entry to the food port. Each chamber was equipped with a syringe pump to deliver 20% sucrose solution in 0.1 ml increments through a stainless-steel tube into one well of the food port and a pellet dispenser to deliver 45-mg food pellets (Bio-Serv, Frenchtown, NJ) into another well of the same port. Both a white noise and tone generator were attached to independent speakers on the wall opposite the levers and food-delivery port. A clicker was also mounted on this wall. A fan mounted to the outer chamber provided ventilation and external noise reduction. A 3-watt, 24-volt house light mounted on the top of the back wall opposite the food port provided illumination, except in Pavlovian blocking experiments for which it was used as a conditioned stimulus. For the Pavlovian blocking behavioral experiment, two stimulus lights were also positioned facing up outside, but immediately adjacent to the chamber at floor level on the front left and back right corners. Chambers used for intracranial self-stimulation contained 2 nose poke ports on the wall with the house light, a smooth plexiglass floor, and rounded wall opposite the nose pokes. They did not contain levers or a food-delivery port. For optogenetic manipulations, chambers were outfitted with an Intensity Division Fiberoptic Rotary Joint (Doric Lenses, Quebec, QC, Canada) connecting the output fiber optic patch cords to a laser (Dragon Lasers, ChangChun, JiLin, China) positioned outside of the chamber.

#### Pavlovian long-delay conditioning

##### Magazine conditioning

Rats first received 2 days of training to learn where to receive the sucrose (20%, 0.1 ml/delivery) and food pellet (45 mg grain; Bio-Serv) outcomes. Each day included 2 sessions, separated by approximately 1 hr, order counterbalanced across days, one with 30 non-contingent deliveries of sucrose and one with 30 grain pellet deliveries (60-s intertrial interval, ITI).

##### Preexposure

To reduce the initial saliency of the auditory stimuli used in subsequent Pavlovian conditioning, subjects received one day of preexposure to the click and white noise stimuli. Click and noise were presented pseudo-randomly for 30 s, 4 times each with a variable 1.5 – 3-min ITI (mean = 2.5 min).

##### Pavlovian conditioning

All rats received 8 sessions of Pavlovian conditioning (1 session/day on consecutive days) to learn to associate each of 2 auditory cues (aka conditioned stimuli; 80-82 db), click (10 Hz) and white noise, with a specific food outcome, sucrose solution or grain pellets. Each 30-s cue terminated with the delivery of its associated outcome. For half the subjects, click terminated in the delivery of sucrose and noise predicted pellets, with the other half receiving the opposite arrangement. Each session consisted of 8 click and 8 white noise presentations. Cues were delivered pseudo-randomly with a variable 1.5 – 3-min ITI (mean = 2.5 min).

#### Pavlovian trace conditioning

##### Magazine conditioning

Rats first received 2 days of training to learn where to receive the chocolate purified pellet (45 mg; Bio-Serv) and unflavored purified pellet (45 mg; Bio-Serv) food rewards. Each day included 2 sessions, separated by approximately 1 hr, order counterbalanced across days, one with 30 non-contingent deliveries of chocolate pellets and one with 30 unflavored pellet deliveries (60-s ITI).

##### Preexposure

To reduce the initial saliency of the auditory stimuli used in subsequent Pavlovian conditioning, subjects received one day of preexposure to the click and white noise stimuli. Click and noise were presented pseudo-randomly for 30-s durations, 4 times each with a variable 1.5 – 3-min ITI (mean = 2.5 min).

##### Pavlovian conditioning

All rats received 8 sessions of Pavlovian conditioning (1 session/day on consecutive days) to learn to associate each of 2 auditory cues (80-82 db), click (10 Hz) and white noise, with a specific food outcome, chocolate or unflavored purified pellets. Each 10-s cue terminated with the delivery of its associated outcome after a 1.5-s trace interval. For half the subjects, click terminated in the delivery of chocolate pellets and noise predicted unflavored pellets, with the other half receiving the opposite arrangement. Each session consisted of 8 click and 8 white noise presentations. Cues were delivered pseudo-randomly with a variable 1 – 4-min ITI (mean = 2.5 min).

##### Unpredicted food reward

Following training, rats received a single session during which they received non-contingent food-pellet deliveries (chocolate or unflavored, counterbalanced; average 60-s ITI, range = 20-110 s). Rats received 5 trials each in which either 1, 2, or 3 food pellets were delivered into the food-delivery port. Trial order was pseudorandomized.

##### Unpredicted aversive airpuff

Rats received a single session in a context different from that of training during which they received 3-8 non-contingent presentations of a single puff of air (Dust Off; Falcon Safety Products, Branchburg NJ) delivered to the top part of the face at on average 35-s ITI.

#### Pavlovian long-delay conditioning with Pavlovian-to-instrumental transfer test

##### Magazine conditioning

Rats first received 2 days of training to learn where to receive the sucrose (20%, 0.1 ml/delivery) and food pellet (45 mg grain; Bio-Serv) rewards. Each day included 2 sessions, separated by approximately 1 hr, order counterbalanced across days, one with 30 non-contingent deliveries of sucrose and one with 30 grain pellet deliveries (60-s ITI).

##### Instrumental conditioning

Rats next received 11 days, minimum, of instrumental conditioning. Each day consisted of 2 training sessions, one with the left lever and one with the right lever, separated by at least 1 hr with order alternated across days. Each action was reinforced with one of the different food outcomes (e.g., left press→grain pellets/right press→sucrose solution). Lever-outcome pairings were counterbalanced at the start of the experiment within each group. Each session terminated after 20 outcomes had been earned or 45 min had elapsed. Actions were continuously reinforced on the first day and then escalated ultimately to a random-ratio (RR) 20 schedule of reinforcement in which a variable number of presses (average = 20) were required to earn a reward.

##### Pavlovian conditioning

All rats received 8 sessions of Pavlovian conditioning (1 session/day on consecutive days) to learn to associate each of 2 auditory cues (80-82 db), click (10 Hz) and white noise, with a specific food outcome, sucrose solution or grain pellets. Each 30-s cue terminated with the delivery of its associated outcome. For half the subjects, click terminated in the delivery of sucrose and noise predicted pellets, with the other half receiving the opposite arrangement. Cue-reward pairings were counterbalanced within groups and with respect to instrumental lever-outcome pairings. Each session consisted of 8 click and 8 white noise presentations. Cues were delivered pseudo-randomly with a variable 1.5 – 3-min ITI (mean = 2.5 min).

##### Instrumental retraining and extinction

Following Pavlovian conditioning, rats received one instrumental retraining session on the RR-20 reinforcement schedule. Rats then received one session of instrumental extinction to establish a low level of pressing. During this single 30-min session both levers were available but pressing was not reinforced.

##### Outcome-specific Pavlovian-to-instrumental transfer tests

Rats next received an outcome-specific Pavlovian-to-instrumental transfer (PIT) test. During the PIT test, both levers were continuously present, but pressing was not reinforced. After 5 min of lever-pressing extinction, each 30-s cue was presented separately 4 times, separated by a fixed 2.5-min ITI, in alternating order. Cue order was counterbalanced across subjects. No outcomes were delivered following cue presentation. Rats next received 2 instrumental retraining sessions. This was followed by 1 Pavlovian retraining session. After retraining, rats were given a second PIT test. This test was identical to the first except the pre-extinction phase was 10 min and each rat received the cues in opposite order to the first test.

#### Pavlovian trace conditioning with outcome-specific devaluation test

##### Magazine conditioning

Rats first received 2 days of training to learn where to receive the chocolate purified pellet (45 mg; Bio-Serv) and unflavored purified pellet (45 mg; Bio-Serv) rewards. Each day included 2 sessions, separated by approximately 1 hr, order counterbalanced across days, one with 30 non-contingent deliveries of chocolate pellets and one with 30 unflavored pellet deliveries (60-s intertrial interval, ITI).

##### Pavlovian conditioning

All rats received 8 sessions of Pavlovian conditioning (1 session/day on consecutive days) to learn to associate each of 2 auditory cues (aka conditioned stimuli; 80-82 db), pulsed tone (1.5 kHz; 2s on/2s off) and white noise, with a specific food outcome, chocolate or unflavored purified pellets. Each 10-s cue terminated with the delivery of its associated outcome after a 1.5-s trace interval. For half the subjects, tone terminated in the delivery of chocolate pellets and noise predicted unflavored pellets, with the other half receiving the opposite arrangement. Each session consisted of 8 tone and 8 white noise presentations. Cues were delivered pseudo-randomly with a variable 1 – 4-min ITI (mean = 2.5 min).

##### Outcome-specific devaluation by conditioned taste aversion

Following training, one of the food rewards was devalued by pairing with the malaise-inducing agent lithium chloride (LiCl). In the conditioning chambers, rats were given 30, non-contingent deliveries of one pellet type (60-s intertrial interval, ITI) followed immediately by a i.p. injection of LiCl (0.3M, 1.5% volume/weight). For the control, rats were given 30, non-contingent deliveries of the other pellet type (60-s intertrial interval, ITI) in the conditioning chamber, without subsequent LiCl injection. Rats received 1 session/day with 16 total sessions (8 devaluation and 8 control) in the order 3 devaluation, 3 control, 3 devaluation, 3 control, 1 devaluation, 2 control, 1 devaluation.

##### Outcome-specific devaluation probe test

24 hr after the last session, rats next received an outcome-specific devaluation probe test. Each 10-s cue was presented separately 8 times, separated by a variable 2.5-min ITI, in alternating order. Cue order was counterbalanced across subjects. No outcomes were delivered following cue presentation.

##### Outcome-specific devaluation consumption test

After the devaluation probe test, rats next received a consumption choice test to confirm the efficacy of the conditioned taste aversion. Rats were given access to 100 pellets of each type in a choice and allowed to consume freely for 20 min.

#### Outcome-specific blocking and Pavlovian-to-instrumental transfer

##### Magazine conditioning

Rats first received 2 days of training to learn where to receive the sucrose (20%, 0.1 ml/delivery) and food pellet (45 mg grain; Bio-Serv) rewards. Each day included 2 sessions, separated by approximately 1 hr, order counterbalanced across days, one with 30 non-contingent deliveries of sucrose and one with 30 grain pellet deliveries (60-s ITI). The house light was off during these sessions.

##### Instrumental conditioning

Rats next received 11 days, minimum, of instrumental conditioning. Each day consisted of 2 separate training sessions, one with the left lever and one with the right lever, separated by at least 1 hr with order alternated across days. Each action was reinforced with one of the different food rewards (e.g., left press→grain pellets/right press→sucrose solution). Lever-outcome pairings were counterbalanced at the start of the experiment within each group. Each session terminated after 20 outcomes had been earned or 45 min had elapsed. Actions were continuously reinforced on the first day and then escalated ultimately to a RR-20 schedule of reinforcement. The house light was off during these sessions.

##### Pavlovian conditioning

Rats received 12 sessions of visual cue Pavlovian conditioning (1 session/day on consecutive days) in a dark operant chamber to learn to associate visual cues with the food outcomes. For rats in the blocking group, each of 2 30-s visual cues, house light or flashing stimulus lights (2 hz), was paired with a specific food outcome, sucrose (20%, 0.1 ml/delivery) or grain pellets (45 mg; Bio-Serv; e.g., house lightꟷsucrose/flashing lightꟷpellet). Cue-reward pairings were counterbalanced within groups and with respect to instrumental lever-outcome pairings. For half the subjects, the house light terminated in the delivery of sucrose and flashing lights predicted pellets, with the other half receiving the opposite arrangement. Each session consisted of 16 house light and 16 flashing light presentations. Cues were delivered pseudo-randomly with a variable 1.5 – 3-min ITI (mean = 2.5 min). Subjects in the control group (behavioral experiment only) were trained to associate a third distinct, 30-s visual stimulus with both food outcomes. Each session consisted of 32 presentations of lights on either side of the outside of the chamber alternating every 2 s (30-s duration; variable 1.5 – 3-min ITI, mean = 2.5 min). On half the trials the 30-s alternating outside lights cue terminated in the delivery of sucrose (20%, 0.1 ml/delivery) and on the other half in in grain pellets (45 mg; Bio-Serv), in pseudorandom order.

##### Instrumental retraining and extinction

Following Pavlovian conditioning, rats received one instrumental retraining session on the RR-20 reinforcement schedule. Rats then received one session of instrumental extinction to establish a low level of pressing. During this single 30-min session both levers were available but pressing was not reinforced.

##### Preexposure

Rats received one day of preexposure to the auditory stimuli. Click and noise were independently presented pseudo-randomly for 30-s, 8 times each with a variable 1.5 – 3-min ITI (mean = 2.5 min).

##### Compound conditioning

Rats next received 4 compound conditioning sessions (1 session/day on consecutive days) in which the house light and flashing stimulus light cues were each presented in compound with a distinct auditory stimulus, click (10 Hz) or white noise (80-82 dB). For half the subjects in each group, the house light was presented simultaneously for 30 s with the click and the flashing lights concurrent noise for 30 s. The other half of subjects received the opposite arrangement. Visual-auditory cue pairings were counterbalanced within groups and with respect to instrumental and visual cue-reward contingencies. For subjects in the blocking group, each compound stimulus terminated in the reward paired with the visual stimulus during initial Pavlovian conditioning (e.g., house light + white noiseꟷsucrose/flashing light + clickerꟷpellet). Compound cue-reward pairings were counterbalanced across subjects in the control group. Each compound conditioning session consisted of 8, 30-s presentations of each compound cue, terminating in the delivery of the associated food outcome. Compound cues were delivered pseudo-randomly with a variable 1.5 – 3-min ITI (mean = 2.5 min).

##### Outcome-specific Pavlovian-to-instrumental transfer tests

Rats next received an outcome-specific PIT test. During the PIT test, both levers were continuously present, but pressing was not reinforced. After 5 min of lever-pressing extinction, each 30-s cue was presented separately 4 times, separated by a fixed 2.5-min ITI, in alternating order. Cue order was counterbalanced across subjects. No outcomes were delivered following cue presentation. The house light was off at test. Rats in the behavioral experiment next received two instrumental retraining sessions, one session of Pavlovian retraining with only visual cue presentations and one day of compound retraining prior to a second PIT test. This test was identical to the first except the pre-extinction phase was 10 min and each rat received the cues in opposite order to the first test.

#### Data collection

Discrete entries and time spent in the food-delivery port and/or lever presses were recorded continuously for each session. For Pavlovian training and PIT test sessions, the 30-s periods prior to each cue onset served as the baseline for comparison of cue-induced changes in lever pressing and/or food-port entries.

### Fiber photometry recordings of dopamine release in the BLA during Pavlovian long-delay conditioning

#### Subjects

Nine male (*N* = 5) and female (*N* = 4) Long Evans rats (Th-cre-littermates, *N* = 6; Charles River Laboratories, *N* = 3) aged 9-11 weeks at the time of surgery were used to record dopamine release in the BLA across Pavlovian long-delay conditioning. Subjects without sufficient fiber photometry GRAB_DA2h_ signal of sufficient quality (*N* = 2) were excluded from the dataset prior to analysis. An additional 3 subjects expressing GFP (2 male) were used to record GFP fluorescence changes as a control during the last Pavlovian conditioning session.

#### Surgery

Rats were infused bilaterally with AAV encoding the GPCR-activation-based dopamine sensor GRAB_DA2h_ (pAAV9-hsyn-GRAB_DA2h, Addgene) or control fluorophore (AAV8-hSYN-GFP). Virus (0.3 µl) was infused bilaterally into the BLA (AP: -2.7; ML: ±5.0; DV: -8.7 males or -8.6 mm females, from bregma). 5 min later, viral injectors were dorsally repositioned in the BLA for a second viral infusion (0.3 µl; DV: -8.4 males or -8.3 mm females). Subjects included in the control experiment received a single viral infusion (0.5 µl; DV: -8.6 mm). Optical fibers (400-µm diameter, 0.37 NA, Neurophotometrics) were implanted bilaterally 0.2 mm dorsal to the first infusion site. Virus was infused at a rate of 0.1 µl/min using 28-gauge injectors and injectors were left in place for 10 min after the second infusion. Experiments commenced approximately 4 weeks after surgery to allow sufficient expression in the BLA.

#### Fiber photometry recordings

Animals were habituated to the optical tether during the magazine conditioning sessions, but no light was delivered. Following magazine training, fiber photometry was used to image GRAB_DA2h_ fluorescent changes in BLA neurons throughout each Pavlovian long-delay conditioning session (*N* = 9) or GFP fluorescence during the last Pavlovian conditioning session (*N* = 3) using a commercial fiber photometry system (Neurophotometrics Ltd.). 470 nm excitation light was adjusted to approximately 80-100 µW at the tip of the patch cord (fiber core diameter: 400 µm; Doric Lenses). Fluorescence emission was passed through a 535 nm bandpass filter and focused onto the complementary metal-oxide semiconductor (CMOS) camera sensor through a tube lens. Samples were collected at 20 Hz using a custom Bonsai^107^ workflow. Time stamps of task events were collected simultaneously through an additional synchronized camera aimed at the Med Associates interface, which sent light pulses coincident with task events. Signals were saved using Bonsai software and exported to MATLAB (MathWorks, Natick, MA) for analysis. Recordings were collected unilaterally from the hemisphere with the strongest fluorescence signal at the start of the experiment.

### Fiber photometry recordings of calcium activity in BLA neurons during Pavlovian long-delay conditioning

#### Subjects

Eight male (*N* = 4) and female (*N* = 4) wildtype rats (Charles River Laboratories, Wilmington, MA) aged approximately 9 weeks at the time of surgery were included in this study to assess BLA neuronal activity changes across Pavlovian long-delay conditioning. No subjects were excluded.

#### Surgery

Rats were infused bilaterally with adeno-associated virus (AAV) expressing the genetically encoded calcium indicator GCaMP6f under control of the calcium/calmodulin-dependent protein kinase (CaMKII) promoter (pENN.AAV5.CAMKII.GCaMP6f.WPRE.SV40, Addgene, Watertown, MA). Virus (0.5 µl) was bilaterally infused into the BLA (AP: -2.9; ML: ± 5.0; DV: -8.8 mm from bregma) at a rate of 0.1 µl/min using 28-gauge injectors. Injectors were left in place for 10 additional min following infusion. Optical fibers (200-µm diameter, 0.37 numerical aperture (NA), Neurophotometrics, San Diego, CA) were implanted bilaterally 0.2 mm dorsal to the infusion site. Experiments commenced approximately 4 weeks after surgery to allow sufficient expression in BLA cell bodies.

#### Fiber photometry recordings

Animals were habituated to the optical tether during the magazine conditioning sessions, but no light was delivered. Following magazine training, fiber photometry was used to image bulk calcium activity in BLA neurons throughout each Pavlovian conditioning session. We simultaneously imaged GCaMP6f and control fluorescence in the BLA using a commercial fiber photometry system (Neurophotometrics Ltd.). Two light-emitting LEDs (470 nm: Ca2+-dependent GCaMP fluorescence; 415 nm: autofluorescence, motion artifact, Ca2+-independent GCaMP fluorescence) were reflected off dichroic mirrors and coupled via a patch cord (fiber core diameter: 200 µm; Doric Lenses, Quebec, Canada) to the implanted optical fiber. The intensity of excitation light was adjusted to ∼80 µW at the tip of the patch cord. Samples were collected at 20 Hz interleaved between the 415 nm and 470 nm excitation channels. Recordings were collected unilaterally from the hemisphere with the strongest fluorescence signal in the 470 nm channel at the start of the experiment.

### Fiber photometry recordings of dopamine release in the BLA during Pavlovian trace conditioning

#### Subjects

10 male (GRAB_DA2h_, *N* = 3; GRAB_DA2m_, *N* = 3) and female (GRAB_DA2h_, *N* =1; GRAB_DA2m_, *N* = 3) Long Evans rats (Th-cre-littermates, *N* = 5; Gad-cre-, *N* = 2; Charles River Laboratories, *N* = 3) aged 7-9 weeks at the time of surgery were used to record dopamine release in the BLA across Pavlovian trace conditioning. Subjects without sufficient quality GRAB_DA_ signal (*N* = 20) were excluded from the dataset prior to analysis. There were no statistical differences in task related signal (AUC) between GRAB_DA2h_ and GRAB_DA2m_ (F_(1, 8)=_1.39, *P* = 0.27) and no interaction between sensor and any other variable of interest (lowest *P*: F_(4, 32)_ = 1.17, *P* = 0.34), so subjects were combined into a single group. An additional 4 (2 male) subjects served as GFP controls.

#### Surgery

Rats were infused bilaterally with AAV encoding GRAB_DA_ (pAAV9-hsyn-GRAB_DA2h, Addgene or AAV9-hsyn-DA4.4, WZ Biosciences) or control fluorophore (AAV8-hSYN-GFP). Virus (0.3 µl) was infused unilaterally into the BLA (AP: -2.7; ML: ±5.0; DV: -8.7 males or -8.6 mm females, from bregma). 5 min later, viral injectors were dorsally repositioned in the BLA for a second viral infusion (0.3 µl; DV: -8.4 males or -8.3 mm females). Optical fibers (400-µm diameter, 0.37 NA, Neurophotometrics) were implanted bilaterally 0.2 mm dorsal to the first infusion site. Virus was infused at a rate of 0.1 µl/min using 28-gauge injectors and injectors were left in place for 10 min after the second infusion. Experiments commenced approximately 4 weeks after surgery to allow sufficient expression in the BLA.

#### Fiber photometry recordings

Animals were habituated to the optical tether during the magazine conditioning sessions, but no light was delivered. Following magazine training, fiber photometry was used to image GRAB_DA_ or GFP fluorescence in BLA neurons throughout each Pavlovian trace conditioning session using a commercial fiber photometry system (Neurophotometrics Ltd.). 470 nm excitation light was adjusted to approximately 80-100 µW at the tip of the patch cord (fiber core diameter: 400 µm; Doric Lenses) and samples were collected at 20 Hz. In a subset of subjects, recordings were made during aversive airpuffs to the face (GRAB_DA2h_: *N* = 2, 2 male; GRAB_DA2m_: *N =* 6, 3 male) and, in a separate session, unpredicted food-pellet reward deliveries (GRAB_DA2h_: *N =* 2, 2 male; GRAB_DA2m_: *N* = 5, 3 male).

### Optogenetic inhibition of VTA_DA_**→**BLA terminals at reward delivery during Pavlovian long-delay conditioning with Pavlovian-to-instrumental transfer test

#### Subjects

Twenty-one male (*N* = 11) and female (*N* = 10) transgenic Th-cre+ (hemizygous) Long Evans rats aged approximately 10 weeks at the time of surgery were used in this study to assess the necessity of VTA_DA_→BLA projection activity for cue-reward learning. Eleven (6 males) served in the experimental group and 10 (5 males) served as controls. Subjects with misplaced optic fibers (*N* = 3) were excluded from the dataset.

#### Surgery

Th-cre rats were randomly assigned to a viral group and infused bilaterally with a cre-dependent AAV encoding either the inhibitory opsin archaerhodopsin T (ArchT; *N* = 11; 6 males; AAV5-CAG-FLEX-ArchT-tdTomato, Addgene) or a tdTomato fluorescent protein control (tdTomato; *N* = 10; 5 males; AAV5-CAG-FLEX-tdTomato, University of North Carolina Vector Core, Chapel Hill, NC). Virus (0.2 µl) was infused bilaterally at a rate of 0.1 µl/min into the VTA (AP: -5.3; ML: ±0.7; DV: -8.3 mm from bregma) using a 28-gauge injector. Injectors were left in place for 10 min following infusion. Optical fibers (200-µm core, 0.39 NA, Thorlabs, Newton, NJ) held in ceramic ferrules (Kientec Systems, Stuart, FL) were implanted bilaterally in the BLA (AP: -2.7; ML: ±5.0; DV: -8.2 mm from bregma). Experiments commenced 4-5 weeks after surgery to allow sufficient expression in VTA_DA_→BLA terminals at the time of manipulation (7-9 weeks post-surgery).

#### Optogenetic inhibition of VTA_DA_**→**BLA projections

Rats received magazine and instrumental training as above. Animals were habituated to the optical tether (200 µm, 0.22 NA, Doric Lenses) for at least the last 2 sessions of instrumental conditioning, but no light was delivered. Optogenetic inhibition was used to attenuate the activity of ArchT-expressing VTA_DA_ axons and terminals in the BLA at the time of cue-reward pairing during each Pavlovian long-delay conditioning session. During each Pavlovian conditioning session, green light (532 nm; 10 mW) was delivered to the BLA via a laser (Dragon Lasers) connected through a ceramic mating sleeve (Thorlabs) to the ferrule implanted on the rat. Light was delivered continuously for 3 s concurrent with each outcome delivery (occurring at cue offset). If the reward was retrieved after the laser had gone off, then the retrieval entry (first food-port entry after reward delivery) triggered an additional 3-s illumination. Light effects were estimated to be restricted to the BLA based on predicted irradiance values (https://web.stanford.edu/group/dlab/cgi-bin/graph/chart.php). Following Pavlovian conditioning, rats proceeded to the PIT tests as described above, during which they were tethered to the optical patch cords, but no light was delivered. The same light delivery procedures were used during Pavlovian retraining in between PIT tests.

### Optogenetic inhibition of VTA_DA_**→**BLA terminals during entire cue-reward period during Pavlovian long-delay conditioning with Pavlovian-to-instrumental transfer test

#### Subjects

Fifteen male (*N* = 8) and female (*N* = 7) transgenic Long Evans rats aged approximately 9 weeks at the time of surgery were used in this study to assess the necessity of VTA_DA_→BLA projection activity for cue-reward learning. Seven (4 males) Th-cre+ (hemizygous) rats served in the experimental group. 1 subject with insufficient viral expression was excluded from the dataset. Eight subjects served in the control group, 4 (2 male) Th-cre+ (hemizygous) rats and 4 (2 male) wildtype Th-cre-littermates. Behavioral data did not differ between the two control types (lowest *P:* F_(1, 6)_ = 1.61, *P* = 0.25) and so they were collapsed into a single control group.

#### Surgery

Th-cre rats were randomly assigned to a viral group and infused bilaterally with a cre-dependent AAV encoding either ArchT (*N* = 7; 4 males; AAV5-CAG-FLEX-ArchT-tdTomato, Addgene) or a tdTomato fluorescent protein control (tdTomato; *N* = 4; 2 males; AAV5-CAG-FLEX-tdTomato). Four (2 male) wildtype Th-cre-littermates were infused bilaterally with the cre-dependent AAV encoding ArchT (AAV5-CAG-FLEX-ArchT-tdTomato, Addgene). Virus (0.2 µl) was infused bilaterally at a rate of 0.1 µl/min into the VTA (AP: -5.3; ML: ±0.7; DV: -8.3 mm from bregma) using a 28-gauge injector. Injectors were left in place for 10 min following infusion. Optical fibers (200-µm core, 0.39 NA, Thorlabs, Newton, NJ) held in ceramic ferrules (Kientec Systems, Stuart, FL) were implanted bilaterally in the BLA (AP: -2.7; ML: ±5.0; DV: -8.2 mm from bregma). Experiments commenced 4-5 weeks after surgery to allow sufficient expression in VTA_DA_→BLA terminals at the time of manipulation (7-9 weeks after surgery).

#### Optogenetic inhibition of VTA_DA_**→**BLA projections

Rats received magazine and instrumental training as above. Animals were habituated to the optical tether (200 µm, 0.22 NA, Doric Lenses) for at least the last 2 sessions of instrumental conditioning, but no light was delivered. Optogenetic inhibition was used to attenuate the activity of ArchT-expressing VTA_DA_ axons and terminals in the BLA throughout cue and reward presentation during each Pavlovian long-delay conditioning session. During each Pavlovian conditioning session, green light (532 nm; 10 mW) was delivered to the BLA via a laser (Dragon Lasers) connected through a ceramic mating sleeve (Thorlabs) to the ferrule implanted on the rat. Light was delivered continuously beginning at the onset of each cue and ending 3 s after cue offset (33 s total). Thus, we inhibited during the entire cue-reward period. If the reward was retrieved after the laser had gone off, then the retrieval entry triggered an additional 3-s illumination. Following Pavlovian conditioning, rats proceeded to the PIT tests as described above, during which they were tethered to the optical patch cords, but no light was delivered. The same light delivery procedures were used during Pavlovian retraining in between PIT tests.

### Optogenetic inhibition of VTA_DA_**→**BLA terminals at reward delivery during Pavlovian trace conditioning with outcome-specific devaluation test

#### Subjects

Twelve male (*N* = 8) and female (*N* = 4) transgenic Long Evans rats aged approximately 10 weeks at the time of surgery were used in this study to assess the necessity of VTA_DA_→BLA projection activity for cue-reward learning. Five (4 males) Th-cre+ (hemizygous) rats served in the experimental group. Subjects that died prior to test (*N* = 2) or with misplaced optic fibers (*N* = 1) were excluded from the dataset. Seven total subjects served in the control group, 4 (3 male) Th-cre+ (hemizygous) rats and 3 (1 male) wildtype Th-cre-littermates. 1 subject that died prior to test was excluded from the dataset. Behavioral data did not differ between the two types of controls (lowest *P*: F_(1, 5)_ = 1.18, *P* = 0.33) and so they were collapsed into a single control group.

#### Surgery

Th-cre rats were randomly assigned to a viral group and infused bilaterally with a cre-dependent AAV encoding either ArchT (*N* = 5; 4 males; AAV5-CAG-FLEX-ArchT-tdTomato, Addgene) or a tdTomato fluorescent protein control (tdTomato; *N* = 4; 3 males; AAV5-CAG-FLEX-tdTomato). Three (1 male) wildtype Th-cre-littermates were infused bilaterally with the cre-dependent AAV encoding ArchT (AAV5-CAG-FLEX-ArchT-tdTomato, Addgene). Virus (0.2 µl) was infused bilaterally at a rate of 0.1 µl/min into the VTA (AP: -5.3; ML: ±0.7; DV: -8.3 mm from bregma) using a 28-gauge injector. Injectors were left in place for 10 min following infusion. Optical fibers (200-µm core, 0.39 NA, Thorlabs, Newton, NJ) held in ceramic ferrules (Kientec Systems, Stuart, FL) were implanted bilaterally in the BLA (AP: -2.7; ML: ±5.0; DV: -8.2 mm from bregma). Experiments commenced 4-5 weeks after surgery to allow sufficient expression in VTA_DA_→BLA terminals at the time of manipulation (7-9 weeks after surgery).

#### Optogenetic inhibition of VTA_DA_**→**BLA projections

Rats received magazine conditioning as above. Animals were habituated to the optical tether (200 µm, 0.22 NA, Doric Lenses) during this training, but no light was delivered. Optogenetic inhibition was used to attenuate the activity of ArchT-expressing VTA_DA_ axons and terminals in the BLA at the time of cue-reward pairing during each Pavlovian trace conditioning session. During each Pavlovian conditioning session, green light (532 nm; 10 mW) was delivered to the BLA via a laser (Dragon Lasers) connected through a ceramic mating sleeve (Thorlabs) to the ferrule implanted on the rat. Light was delivered continuously for 3 s concurrent with each reward delivery (occurring 1.5 s after cue offset). If the outcome was retrieved after the laser had gone off, then the retrieval entry triggered an additional 3-s illumination. Thus, we inhibited at each reward delivery, without inhibiting at cue offset. Rats received 5 conditioning sessions to avoid negatively deflecting VTA_DA_→BLA activity once BLA dopamine reward responses attenuate with learning. Following Pavlovian conditioning, rats proceeded to the outcome-specific devaluation and probe test as described above. Rats were tethered to the optical patch cords during the probe test, but no light was delivered.

### Outcome-specific Pavlovian blocking and Pavlovian-to-instrumental transfer

#### Subjects

Thirty-two male (*N* = 22) and female (*N* = 10) Long Evans rats (Charles River) aged approximately 8 weeks at the start of the experiment were used in this study to evaluate the extent to which previously learned cues could block the encoding of novel identity-specific cue-reward memories. Prior to the start of behavioral training, subjects were randomly assigned to Blocking (*N* = 16, 11 male) or Control (*N* = 16, 11 male) groups. Rat were trained and tested using the Outcome-specific blocking and Pavlovian-to-instrumental transfer procedures described above.

### Optical stimulation of VTA_DA_**→**BLA terminals at reward delivery during Pavlovian blocking with Pavlovian-to-instrumental transfer

#### Subjects

Twenty-four male (*N* = 12) and female (*N* = 12) transgenic TH-cre+ (hemizygous) Long Evans rats aged between 9-12 weeks at the time of surgery were used in this study. Subjects with misplaced optical fibers (*N* = 2) or lacking viral expression (*N* = 2) were excluded from the dataset.

#### Surgery

Th-cre rats were randomly assigned to a viral group and infused bilaterally with a cre-dependent AAV encoding either the excitatory opsin channelrhodopsin (ChR2; *N* = 11, 6 male; AAV5-EF1a-DIO-hChR2(H134R)-eYFP, University of North Carolina Vector Core) or an enhanced yellow fluorescent protein control (eYFP; *N* = 13, 6 males; pAAV5-Ef1a-DIO-eYFP, Addgene). Virus (0.2 µl) was infused bilaterally at a rate of 0.1 µl/min into the VTA (AP: -5.3; ML: ±0.7; DV: -8.3 mm from bregma) using a 28-gauge injector. Injectors were left in place for 10 min following viral infusions. Optical fibers (200 µm core, 0.39 NA, Thorlabs) held in ceramic ferrules (Kientec Systems) were implanted bilaterally in the BLA (AP: -2.7; ML: ±5.0; DV: -8.2 mm from bregma). Experiments commenced approximately 2 weeks after surgery to allow sufficient expression in VTA_DA_→BLA axon terminals at the time of optical manipulation (7-8 weeks after surgery).

#### Optogenetic stimulation of VTA_DA_**→**BLA projections

Rats received magazine conditioning, instrumental training, and visual cue Pavlovian conditioning as described for the Outcome-specific blocking and Pavlovian-to-instrumental transfer procedures above. All subjects received the blocking condition. Animals were habituated to the optical tether (200 µm, 0.22 NA, Doric Lenses) for at least the last 2 sessions of instrumental conditioning and the last two days of visual cue Pavlovian conditioning, but no light was delivered. Optogenetic excitation was used to stimulate the activity of ChR2-expressing VTA_DA_ axons and terminals in the BLA at the time of each cue-reward pairing during each compound conditioning session. During each compound conditioning session, blue light (473 nm; 10 mW; 25-ms pulse width) was delivered to the BLA via a laser (Dragon Lasers) for 3 s at a rate of 20 Hz concurrent with each reward delivery. We selected this stimulation frequency to match the upper end firing rate of VTA_DA_ neurons detected in response to reward^3, 54^ similar to prior work on the VTA→BLA pathway^108^. Following compound conditioning, rats proceeded to the PIT test as described above, during which they were tethered to the optical patch cords, but no light was delivered.

#### Intracranial self-stimulation

Following the PIT test, rats received 2 sessions (1 session/day) of intracranial self-stimulation (ICSS) testing. This occurred in a distinct context from the prior conditioning and testing. This context had a smooth plexiglass rather than grid floor, round right-side wall and no levers or food-delivery port. During each 1-hr session animals were allowed to nose poke in 2 ports positioned on the left and right side of the left wall of the operant chamber. Nose pokes into the active port triggered 1-s blue light (473nm; 10 mW; 25-ms pulse width; 20 Hz) delivery to the BLA. Subsequent nose pokes during the 1-s light-delivery period were recorded but did not extend light delivery. Inactive port pokes were also recorded. For half of the subjects in each group, the left port was active and the right inactive, with the opposite arrangement for the other half.

### Histology

Following behavioral experiments, rats were deeply anesthetized with Nembutal and transcardially perfused with phosphate buffered saline (PBS) followed by 4% paraformaldehyde (PFA). Brains were removed and post-fixed in 4% PFA overnight, placed into 30% sucrose solution, then sectioned into 30-μm slices using a cryostat and stored in cryoprotectant. Slices were rinsed in a DAPI solution for 4 min (5 mg/mL stock, 1:10000), washed 3 times in PBS for 15 min, mounted on slides and coverslipped with ProLong Gold mounting medium. Images were acquired using a Keyence BZ-X710 microscope (Keyence, El Segundo, CA) with a 4x, 10x, and 20x objective (CFI Plan Apo), CCD camera, and BZ-X Analyze software.

GFP fluorescence was used to confirm expression of GCaMP6f in BLA cell bodies. Immunofluorescence was used to confirm expression of GRAB_DA_ in the BLA. Floating coronal sections were washed 3 times in 1x PBS for 30 min and then blocked for 1–1.5 hr at room temperature in a solution of 3% normal goat serum and 0.3% Triton X-100 dissolved in PBS. Sections were then washed 3 times in PBS for 15 min and incubated in blocking solution containing chicken anti-GFP polyclonal antibody (1:1000; Abcam, Cambridge, MA) with gentle agitation at 4°C for 18–22 hr. Sections were next rinsed 3 times in PBS for 30 min and incubated with goat anti-chicken IgY, Alexa Fluor 488 conjugate (1:500; Abcam) in blocking solution at room temperature for 2 hr. Sections were washed a final 2 times in PBS for 10 min.

tdTomato fluorescence with a Th costain was used to confirm expression of ArchT-tdTomato in VTA_DA_ neurons. Floating coronal sections were washed 3 times in 1x PBS for 30 min and then blocked for 2 hr at room temperature in a solution of 3% normal donkey serum and 0.2% Triton X-100 dissolved in PBS. Sections were then washed 3 times in PBS for 15 min and incubated in blocking solution containing rabbit anti-TH antibody (1:1000; EMD Millipore, Burlington, MA) with gentle agitation at 4°C for 44-48 hr. Sections were next rinsed 3 times in PBS for 30 min and incubated with goat anti-rabbit IgG, Alexa Fluor 488 conjugate (1:500; Thermofisher Scientific, Waltham, MA) in blocking solution at room temperature for 2 hr. Sections were washed a final 2 times in PBS for 10 min. Immunofluorescence was also used to confirm expression of ArchT-tdTomato in axons and terminals in the BLA. Floating coronal sections were washed 2 times in 1x PBS for 10 min and then blocked for 2 hr at room temperature in a solution of 10% normal goat serum and 0.5% Triton X-100 dissolved in PBS. Sections were then washed 3 times in PBS for 15 min and incubated in blocking solution containing rabbit anti DsRed polyclonal antibody (1:1000; EMD Millipore, Burlington, MA) with gentle agitation at 4°C for 18-22 hr. Sections were next rinsed 3 times in blocking solution for 30 min and incubated with goat anti-rabbit IgG, Alexa Fluor 594 conjugate (1:500; Thermofisher Scientific) in blocking solution at room temperature for 2 hr. Sections were washed a final 2 times in PBS for 10 min.

eYFP fluorescence with a Th costain was used to confirm expression of ChR2-eYFP expression in VTA_DA_ neurons. Staining procedures were as described above using a secondary goat anti-rabbit Alexa 594 antibody (Thermofisher Scientific). Immunofluorescence following procedures described for GFP amplification also described above were used to confirm expression of ChR2 in axons and terminals in the BLA.

### Data analysis

#### Behavioral analysis

Behavioral data were processed with Microsoft Excel (Microsoft, Redmond, WA). Press rates on the last 2 sessions of instrumental training were averaged across levers then across days and compared between groups to test for any pre-existing group differences in instrumental behavior. For Pavlovian long-delay conditioning, conditional food-port approach responses during the Pavlovian and compound conditioning sessions were assessed by comparing the rate of entries into the food-delivery port (entries/min) during the 30-s cue periods relative to the 30-s baseline periods prior to cue onset (preCue). Because cue periods were shorter, for trace conditioning, Pavlovian conditional food-port approach responses during the Pavlovian conditioning sessions were assessed by comparing the percentage of time spent in the food-delivery port during the 10-s cue periods relative to the 10-s baseline preCue periods. Data were averaged across trials for each cue and then averaged across the two cues. For PIT tests, entry rate into the food-port during the 30-s cues were also compared to the baseline 30-s preCue periods. Data were averaged across trials for each cue and then averaged across cues. Lever press rates (presses/min) during the 30-s baseline preCue periods were compared to that during the 30-s cue periods to capture the cue-induced change in lever pressing. Lever presses were separated for presses on the lever that, during training, earned the same outcome as the upcoming or presented cue (Same presses) versus those on the other available lever (Different presses). Data was separated into Same v. Different presses for each preCue and cue period, averaged across trials, then averaged across cue types. To account for baseline press rates and evaluate the selective cue-induced change in lever pressing, we computed an elevation ratio for each lever [(Cue:Same presses)/(Cue:Same presses + preCue:Same presses)] and [(Cue:Different presses)/(Cue:Different presses + preCue:Different presses)]. When two PIT tests were conducted, food-port entry rate, lever-press rates, and elevation ratios were averaged across PIT tests. For devaluation tests, percent time in the food-delivery port was averaged across the 10-s baseline preCue periods and compared with that during the 10-s cue periods, separated by the cue that signaled the non-devalued outcome v. that signaling the currently devalued (i.e., pre-fed) outcome. Elevation ratios were computed for each cue [(Cue: non-Devalued %time)/(Cue: non-Devalued %time + preCue %time)] and [(Cue: Devalued %time)/(Cue: Devalued %time + preCue %time)]. For ICSS sessions, the total number of nose pokes into the active and active ports were compared across the two sessions.

#### GRAB_DA_ fiber photometry analysis

Data were pre-processed using a custom-written pipeline in MATLAB (MathWorks). The 470 nm signal was resampled to 19.5 Hz and then was divided by a second-order exponential fitted to the raw data to account for attenuation in fluorescence resulting from photobleaching across the session. The resampled and fitted data were then Z-scored. Area under the curve (AUC) was calculated for each individual aligned trace within each session using a trapezoidal function. For the Pavlovian long-delay conditioning, we used 2-s preCue baseline, cue onset, and Cue offset/reward delivery windows. To match the duration of the trace interval, for Pavlovian trace conditioning, we used 1.5-s preCue baseline, Cue onset, Cue offset, and reward delivery windows. We compared data across conditioning sessions 1, 2, 3/4, 5/6, and 7/8. Thus, data from the mid and latter training sessions were averaged across 2-session bins. All subjects had reliable data from at least one session per bin. Session data were excluded if artifactual signal due to excessive motion or patch cord twisting was detected for at least half of the trials (Long-delay conditioning: *N* = 3 sessions from *N* = 2 subjects; Trace conditioning: *N* = 1 sessions from *N* = 2 subjects). Two subjects without reliable data from at least one session per bin were excluded from the long-delay conditioning experiment, and two subjects were excluded from the trace conditioning experiment. We were able to obtain reliable imaging data from all 8 training sessions from *N* = 7/9 subjects for the long-delay conditioning experiment and from *N* = 8/10 subjects for the trace conditioning experiment (Figure 2-1). When evaluating the data from subjects for which we collected reliable recordings from each training session, there were no significant statistical interaction between Session and Events of interest across the data within each bin (Long-delay conditioning: lowest *P*: F_(1.11, 6.68)_ = 1.66, *P* = 0.25; Trace conditioning: lowest *P*: F_(1.68, 11.73)_ = 0.97, *P* = 0.39), justifying the collapse across training bins.

#### GCaMP6f fiber photometry analysis

Data were pre-processed using a custom-written pipeline in MATLAB (MathWorks, Natick, MA). Using least-squares linear regression, the 415 nm signal was fit to the 470 nm signal. Change in fluorescence (ΔF/F) at each time point was calculated by subtracting the fitted 415 nm signal from the 470 nm signal and normalizing to the fitted 415 nm data [(470-fitted 415)/fitted 415)]. The ΔF/F data were resampled to 19.5 Hz then Z-scored [(ΔF/F - mean ΔF/F)/std(ΔF/F)]. Using a custom MATLAB workflow, Z-scored traces were then aligned to cue onset for each trial. Peak magnitude was calculated on the Z-scored trace for each trial using 5-s preCue baseline, cue onset, and postCue offset/outcome delivery windows. Data were averaged across trials and then across cues. Session data were excluded if no transient calcium fluctuations were detected on the 470 nm channel above the isosbestic channel or if poor linear fit was detected due to excessive motion artifact (*N* = 2 sessions from *N* = 2 subjects). To examine the progression in BLA activity across training, we compared data across conditioning sessions 1, 2, 3/4, 5/6, and 7/8. Thus, data from the mid and latter training sessions were averaged across 2-session bins. All subjects had reliable data from at least one session per bin.

#### Statistical analysis

Datasets were analyzed by two-tailed, paired and unpaired Student’s *t* tests, two-, or three-way repeated-measures analysis of variance (ANOVA), and simple linear regression as appropriate (GraphPad Prism, GraphPad, San Diego, CA; SPSS, IBM, Chicago, IL). For the few datasets that were slightly non-normal, results were cross-checked using non-parametric statistics and the findings were identical. We opted to use parametric statistics for consistency across experiments and given evidence that ANOVA is robust to slight non-normality^109, 110^. For well-established behavioral effects (PIT), multiple pairwise comparisons were used for *a priori post hoc* comparisons based on a logical extension of Fisher’s protected least significant difference procedure for controlling familywise Type I error rates^111^. All other *post hoc* tests were corrected for multiple comparisons using the Bonferroni method and used to clarify main and interaction effects. Greenhouse-Geisser correction was applied to mitigate the influence of unequal variance between conditions. Alpha levels were set at *P* < 0.05.

### Sex as a biological variable

Male and female rats were used in approximately equal numbers for each experiment, but the *N* per sex was underpowered to examine sex differences. Sex was therefore not included as a factor in statistical analyses, though individual data points are visually disaggregated by sex.

### Rigor and reproducibility

Group sizes were estimated *a priori* based on prior work using male Long Evans rats in this behavioral task^106, 112, 113^ and to ensure counterbalancing of Cue-reward and Lever-outcome pairings. Investigators were not blinded to viral group because they were required to administer virus. All behaviors were scored using automated software (MedPC). Each experiment included at least 1 replication cohort and cohorts were balanced by viral group, Cue-reward and Lever-reward pairings, hemisphere etc. prior to the start of the experiment.

### Data and code availability

All data that support the findings of this study is available in the supplemental files. Custom-written MATLAB code is available from the corresponding author upon request and the basic code is available via Dryad (https://doi.org/10.5068/D1109S).

## SUPPLEMENTAL MATERIALS

**Supplemental Figure 1-1.**
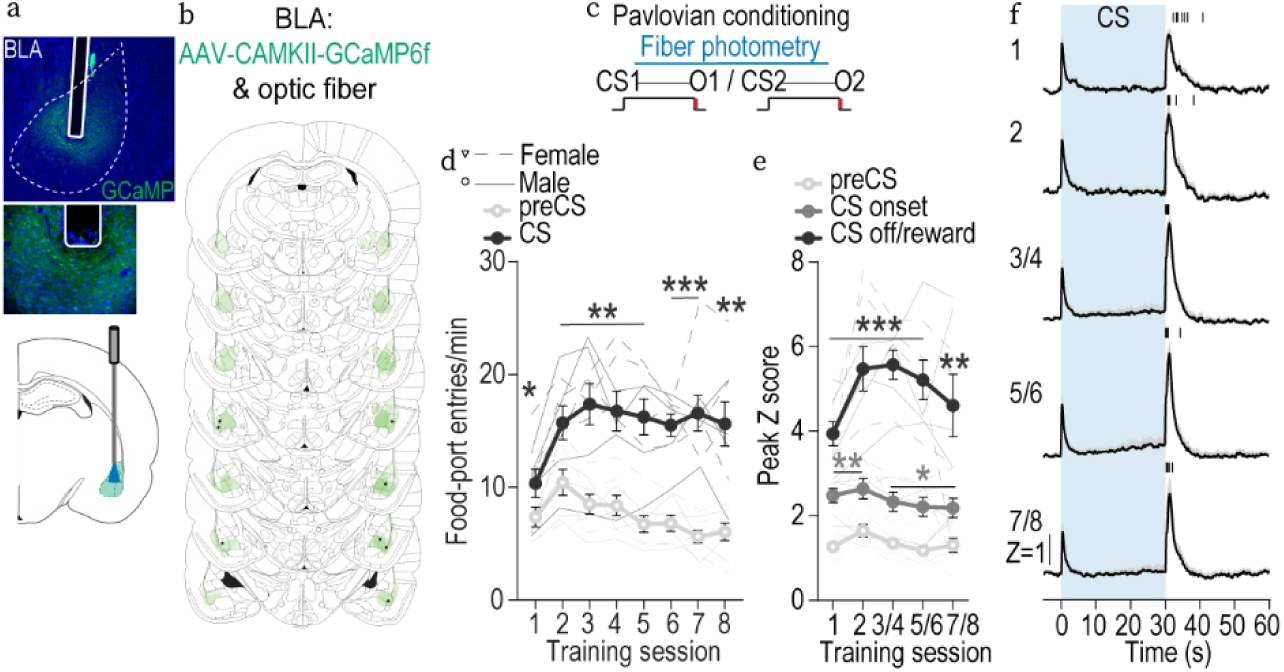
BLA neurons are active during cue-reward encoding. To characterize the endogenous activity of BLA neurons, we used fiber photometry to record fluorescent activity of the genetically encoded calcium indicator GCaMP6f^114^ in the BLA of male and female rats. **(a)** Top: Representative fluorescent image of GCaMP6f expression and fiber placement in the BLA. Bottom: Schematic of fiber photometry approach for bulk calcium imaging in BLA neurons. **(b)** Schematic representation of GCaMP6f expression and placement of optical fiber tips in BLA for all subjects. **(c)** Pavlovian long-delay conditioning procedure schematic. CS, 30-s conditioned stimulus (aka, “cue”, white noise or click) followed immediately by reward outcome (O, sucrose solution or grain pellet). **(d)** Food-port entry rate during the cue relative to the preCue baseline period, averaged across trials and across the 2 cues for each Pavlovian conditioning session. Thin lines represent individual subjects. Across training, rats developed a Pavlovian conditional approach response of entering the food-delivery port during cue presentation. Training x Cue: *F*_(2.44, 17.07)_ = 7.97, *P =* 0.002; Training: *F*_(3.30, 23.10)_ = 4.85, *P =* 0.008; Cue: *F*_(1, 7)_ = 80.33, *P* < 0.0001. \**P* < 0.05, \*\**P* < 0.01, \*\*\**P* < 0.001 relative to preCue, Bonferroni correction. **(e-f)** BLA neurons are active during the encoding of cue-reward memories. BLA neurons were robustly activated both at cue onset and offset when the outcome was delivered. Cue onset responses beginning on the first conditioning sessions have been detected previously^36^. These novelty responses rapidly attenuate if the stimuli are not associated with reward^36^. **(e)** Trial-averaged quantification of maximal (peak) GCaMP6f Z-score ΔF/F during the 5-s period following cue onset or outcome delivery compared to the equivalent baseline period immediately prior to cue onset. Training x Event: *F*_(2.52, 17.61)_ = 3.94*, P =* 0.03; Event: *F*_(1.39, 9.71)_ = 58.63, *P* < 0.0001; Training *F*_(1.71, 11.97)_ = 2.30, *P =* 0.15. \**P* < 0.05, \*\**P* < 0.01, \*\*\**P* < 0.001 relative to preCue baseline, Bonferroni correction. **(f)** Trial-averaged GCaMP6f fluorescence changes (Z-score ΔF/F) in response to cue presentation (blue) and outcome delivery across days of training. Shading reflects between-subjects s.e.m. Tick marks represent time of outcome collection for each subject. Data from the last six sessions were averaged across 2-session bins (3/4, 5/6, and 7/8). *N* = 8, 4 male. Consistent with prior evidence^36^, BLA neurons are activated by rewards and their predictors. BLA activation is particularly robust when the cues can become linked to the identifying features of the rewards they predict. Although these data likely reflect both somatic and non-somatic calcium activity, they are consistent with prior electrophysiological evidence that BLA neurons respond to reward during learning^56–59^.

**Supplemental Figure 1-2.**
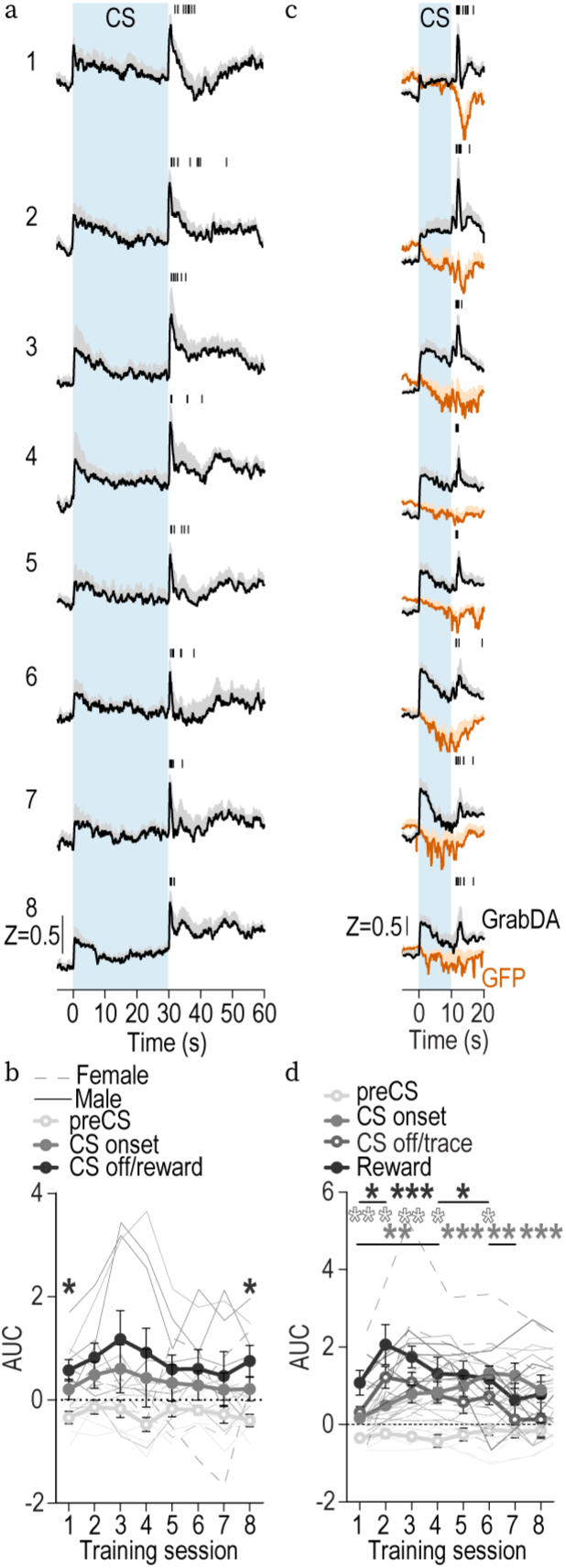
Dopamine release in BLA during cue-reward learning across each of the 8 Pavlovian conditioning sessions. **(a)** Trial-averaged GRAB_DA2h_ fluorescence changes (Z-score) in response to cue presentation (blue) and reward delivery across each of the 8 Pavlovian conditioning sessions. **(b)** Trial-averaged quantification of BLA GRAB_DA_ Z-scored signal AUC during the 2-s period following cue onset or reward delivery compared to the equivalent baseline period immediately prior to cue onset. Event: F_(1.85, 11.07)_ = 4.90, *P* = 0.03; Training: F_(2.34, 14.03)_ = 1.13, *P* = 0.36; Training x Event: F_(3.45, 20.99)_ = 0.59, P=0.65. \**P* < 0.05, relative to preCue baseline, Bonferroni correction. *N* = 7, 4 male. **(c)** Trial-averaged GRAB_DA_ fluorescence changes (Z-score) in response to cue presentation and reward delivery across each of the 8 Pavlovian conditioning sessions. **(d)**Trial-averaged quantification of BLA GRAB_DA_ Z-scored signal AUC during the 1.5-s period following cue onset, cue offset (trace interval), or reward delivery compared to the equivalent baseline period immediately prior to cue onset. Event: F_(2.06, 14.40)_ = 13.24, *P* = 0.0005; Training: F_(3.62, 25.33)_ = 2.43, *P* = 0.08; Training x Event: F_(3.60, 25.17)_ = 2.60, *P* = 0.07. \**P* < 0.05, *\*\*P < 0.01,* \*\*\**P* < 0.001, relative to preCue baseline, Bonferroni correction. (GRAB_DA2h_: *N* = 3, 2 male; GRAB_DA2m_: *N* = 5, 3 male). The slope of the BLA dopamine reward response across training was significantly negative (β = -0.13, confidence interval -0.25 – -0.007; F_(1,62)_ = 4.49, *P* = 0.04) and signifantly different (F_(1,124)_ = 13.33, *P* = 0.0004) from the slope of the BLA dopamine cue-onset response across training, which was significantly positive (β = 0.13, confidence interval 0.06 – 0.20; F_(1,62)_ = 13.53, *P* = 0.0005).

**Supplemental Figure 1-3:**
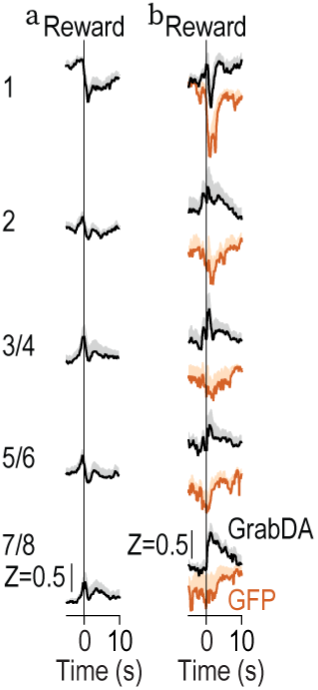
GRAB_DA_ responses to reward collection. **(a)** Trial-averaged GRAB_DA2h_ fluorescence changes (Z-score) in response to reward collection across Pavlovian long-delay conditioning. Shading reflects between-subjects s.e.m. Data from the last six sessions were averaged across 2-session bins (3/4, 5/6, and 7/8). *N* = 9, 5 male. **(b)** Trial-averaged GRAB_DA_ fluorescence changes (Z-score) in response to reward collection across Pavlovian trace conditioning. Shading reflects between-subjects s.e.m. Data from the last six sessions were averaged across 2-session bins (3/4, 5/6, and 7/8). GRAB_DA2h_: *N* = 4, 3 male; GRAB_DA2m_: *N* = 6, 3 male.

**Supplemental Figure 1-4:**
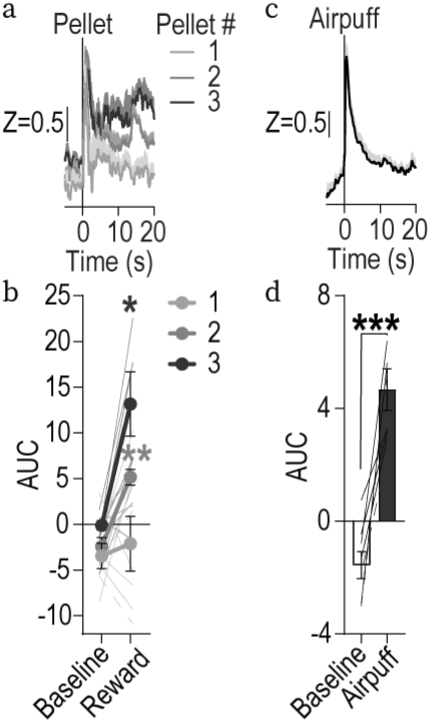
GRAB_DA_ responses to unpredicted rewarding and aversive events. **(a)** Trial-averaged GRAB_DA_ fluorescence changes (Z-score) in response to unpredicted delivery of 1, 2, or 3 food pellets. **(b)** Trial-averaged quantification of BLA GRAB_DA_ Z-scored signal AUC during the 20-s period following pellet delivery. Reward period x Magnitude: F_(1.92, 11.50)_ = 12.46, *P* = 0.001; Magnitude: F_(1.94, 11.66)_ = 11.04, *P* = 0.002; Reward: F_(1, 6)_ = 7.86, *P* =0.03. GRAB_DA2h_ : *N =* 2, 2 male; GRAB_DA2m_ : *N* = 5, 3 male **(c)** Trial-averaged GRAB_DA_ fluorescence changes (Z-score) in response to unpredicted puff of air to the face. **(d)** Trial-averaged quantification of BLA GRAB_DA_ Z-scored trace AUC during the 5-s period following airpuff delivery relative to 5-s preAirpuff baseline. t_(7)_ = 5.88, *P* = 0.0006. GRAB_DA2h_: *N* = 2, 2 male; GRAB_DA2m_: *N =* 6, 3 male.

**Supplemental Figure 2-1.**
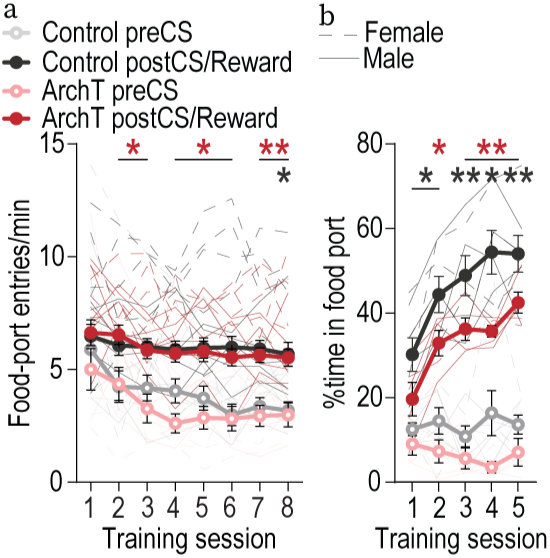
Inhibition of VTA_DA_→BLA projections does not disrupt reward collection during Pavlovian conditioning. There was no effect of optical inhibition of VTA_DA_→BLA projections at reward delivery on collection of the food outcomes. **(a)** Entries into the food-delivery port during the 30-s periods before and after cue presentation during Pavlovian long-delay conditioning. Rats entered the food-delivery port during the 30-s postcue/reward-delivery period more than the preCue baseline period and similarly between groups. Training x Period: F_(4.94,93.85)_ = 3.00, *P =* 0.02; Training: F_(3.13, 59.48)_ = 8.51, *P <* 0.0001; Period: F_(1,19)_ = 72.60, *P <* 0.0001; Virus: F_(1,19)_ = 0.47, *P =* 0.50; Training x Virus: F_(7,133)_ = 0.65, *P =* 0.72; Virus x Period: F_(1,19)_ = 0.87, *P =* 0.36; Training x Virus x Period: F_(7,133)_ = 0.71, *P =* 0.66. ArchT, *N* = 11, 6 males; tdTomato, *N* = 10, 5 males. **(b)** Percent time spent in the food-delivery port during the 10-s preCue baseline and 10-s postCue offset (including trace interval and reward delivery period) periods during Pavlovian trace conditioning. Rats entered the food-delivery port during the 10-s postCue period more than the preCue period and similarly between groups. Training x Period: F_(1.93,19.27)_ = 9.68, *P =* 0.001; Training: F_(2.59, 25.88)_ = 9.28, *P =* 0.0004; Period: F_(1,10)_ = 138.50, *P <* 0.0001; Virus: F_(1,10)_ = 14.94, *P =* 0.003; Training x Virus: F_(4, 40)_ = 1.35, *P =* 0.27; Virus x Period: F_(1,10)_ = 1.37, *P =* 0.27; Training x Virus x Period: F_(4, 40)_ = 0.05, *P =* 0.996. ArchT, *N* = 5, 4 males; Control, *N* = 7, 4 males (3 WT/cre-dependent ArchT; 4 Th-cre/cre-dependent tdTomato).

**Supplemental Figure 2-2.**
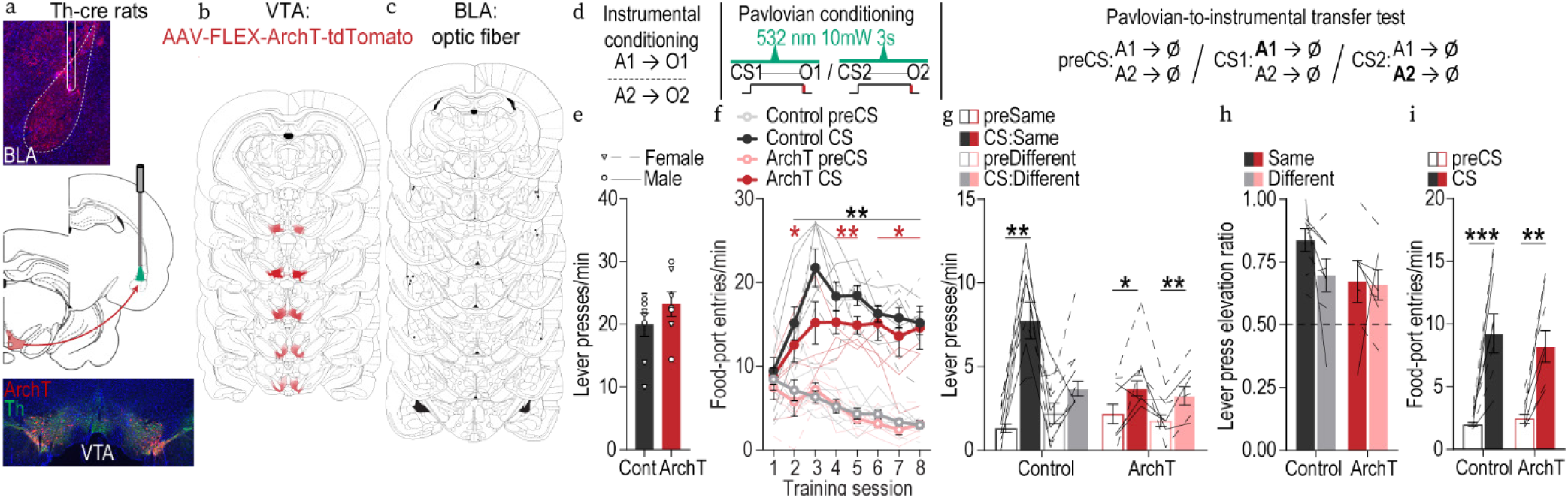
Optical inhibition of VTA_DA_→BLA projections throughout cue and reward during learning attenuates the encoding of identity-specific cue-reward memories. We cre-dependently expressed ArchT bilaterally in VTA_DA_ neurons of male and female Th-cre rats and implanted optical fibers bilaterally over BLA. **(a)** Bottom: Representative fluorescent image of cre-dependent ArchT-tdTomato expression in VTA cell bodies with coexpression of Th in Th-Cre rats. Middle: Schematic of optogenetic strategy for bilateral inhibition of VTA_DA_ axons and terminals in the BLA of Th-cre rats. Top: Representative image of fiber placement in the vicinity of immunofluorescent ArchT-tdTomato-expressing VTA_DA_ axons and terminals in the BLA. **(b)** Schematic representation of cre-dependent ArchT-tdTomato expression in VTA and **(c)** placement of optical fiber tips in BLA for all subjects. For half of the control group, we expressed cre-dependent tdTomato in the VTA of Th-cre male and female rats. For the other half, wildtype rats were infused with cre-dependent ArchT (which did not express owing to the lack of cre recombinase) into the VTA. Both groups received bilateral optical fibers above the BLA. Thus, we control for light delivery, viral expression, and genotype. There were no significant behavioral differences between each type of control (lowest *P:* F_(1, 6)_ = 1.61, *P* = 0.25). **(d)** Procedure schematic. A, action (left or right lever press); CS, 30-s conditioned stimulus (aka, “cue”, white noise or click) followed immediately by reward outcome (O, sucrose solution or grain pellet). **(e)** Rats first received 11 sessions of instrumental conditioning, without manipulation, in which one of two different lever-press actions each earned one of two distinct food rewards (e.g., left press→sucrose/right press→pellets). Lever-press rate averaged across levers and across the final 2 instrumental conditioning sessions. t_(13)_ = 1.20, *P =* 0.25. Data points represent individual subjects. **(f)** Rats then received Pavlovian conditioning. During each of the 8 Pavlovian conditioning sessions, each of 2 distinct, 30-s, auditory cues was presented 8 times and terminated in the delivery of one of the food rewards (e.g., white noiseꟷsucrose/clickꟷpellets). VTA_DA_→BLA projections were optically inhibited (532 nm, 10 mW, 33 s) during the entirety of each cue-reward period. Light turned on at the onset of each cue and off 3 s following reward delivery. Optical inhibition of VTA_DA_→BLA projections through the cue and reward period did not disrupt development of a Pavlovian conditional goal-approach response. Food-port entry rate during the cue relative to the preCue baseline period, averaged across trials and across the 2 cues for each Pavlovian conditioning session. Thin lines represent individual subjects. Training x CS: F_(3.30, 42.87)_ = 20.69, *P <* 0.0001; CS: *F*_(1, 13)_ = 295.60, *P <* 0.0001; Training: *F*_(3.03.,39.42)_ = 4.13, *P =* 0.01; Virus: F_(1,13)_ = 1.61, *P =* 0.23; Training x Virus: F_(7,91)_ = 0.37, *P =* 0.92; Virus x Cue: F_(1,13)_ = 3.05, *P* = 0.10; Training x Virus x CS: F_(7,91)_ = 2.17, *P* = 0.04. By the end of training both groups showed similar elevation in food-port approach during the cues. **(g-i)** We next gave subjects an outcome-specific Pavlovian-to-instrumental transfer (PIT) test, without manipulation. Controls learned the identity-specific cue-reward memories as evidenced by their ability to use the cues to selectively elevate pressing on the lever associated with the same outcome as predicted by the cue. Conversely, the cues were not capable of guiding lever-press choice in the group for which VTA_DA_→BLA projections were inhibited during Pavlovian conditioning. Rather, for these subjects, the cues caused a general increase in pressing across both levers. **(g)** Trial-averaged lever-press rates during the preCue baseline periods compared to press rates during the cue periods separated for presses on the lever that, in training, delivered the same outcome as predicted by the cue (Same) and pressing on the other available lever (Different). Virus x Lever x Cue: F_(1, 13)_ = 7.35, *P* =0.02; Virus: F_(1, 13)_ = 4.59, *P =* 0.05; Lever: F_(1, 13)_ = 5.76, *P =* 0.03; Cue: F_(1, 13)_ = 58.87, *P <* 0.0001; Virus x Lever: F_(1, 13)_ = 1.91, *P =* 0.19; Virus x Cue: F_(1, 13)_ = 12.00, *P* = 0.004; Lever x Cue period: F_(1, 13)_ = 7.56, *P =* 0.02. Inhibition of VTA_DA_→BLA projections during cue-reward learning prevents subjects from learning identity-specific cue-reward memories, but does not prevent the assignment of general incentive properties to the cues that supports non-discriminate cue-induced motivation. **(h)** Elevation in lever presses on the Same lever [(Same lever presses during cue)/(Same presses during cue + Same presses during preCue)], relative to the elevation in pressing on the Different lever [(Different lever presses during cue)/(Different presses during cue + Different presses during preCue)], averaged across trials and across cues during the PIT test. Virus: F_(1, 13)_ = 2.21, *P =* 0.16; Lever: F_(1, 13)_ = 1.67, *P =* 0.22; Virus x Lever: F_(1, 13)_ = 1.14, *P =* 0.30. Lines represent individual subjects. **(i)** As in training, during the PIT test the conditional goal-approach response was similar between groups, further indicating that even longer duration inhibition of VTA_DA_→BLA projections during cue-reward learning does not disrupt development of conditional responses. Food-port entry rate during the cues relative to the preCue baseline periods, averaged across trials and across the 2 cues during the PIT test. Cue: F_(1, 13)_ = 44.71, *P* < 0.0001; Virus: F_(1, 13)_ = 0.08, *P =* 0.79; Virus x Cue: F_(1, 13)_ = 0.61, *P =* 0.45. \**P* < 0.05, \*\**P* < 0.01, \*\*\**P* < 0.001, Bonferroni correction. ArchT, *N* = 7, 4 males; Control *N* = 8, 4 Th-cre/tdTomato 2 males, 4 wildtype cre-dependent ArchT 2 males. These data confirm that VTA_DA_→BLA projections are needed to link the identifying details of the reward to a predictive cue, but not to reinforce a conditional response or to assign general incentive properties to the cue to support general motivation.

**Supplemental Figure 4-1.**
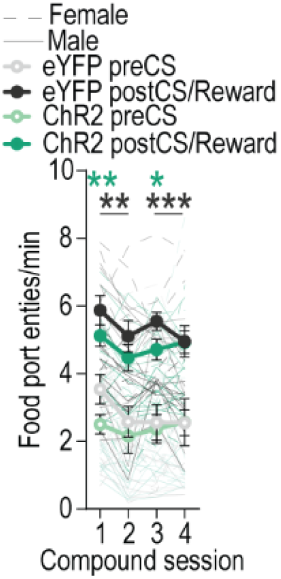
Stimulation of VTA_DA_→BLA projections does not affect reward collection during compound conditioning. There was no effect of optical stimulation of VTA_DA_→BLA projections paired with reward delivery on collection of the food outcomes. Rats entered the food-delivery port during the 30-s postCue/reward-delivery period more than the preCue baseline period and similarly between groups. Period: F_(1, 22)_ = 46.80, *P <* 0.0001; Training: F_(1.50, 32.90)_ = 3.70, *P =* 0.047; Virus: F_(1, 22)_ = 1.89, *P =* 0.18; Training x Virus: F_(3, 66)_ = 1.48, *P =* 0.23; Training x Period: F_(2.55, 56.04)_ = 0.22, *P =* 0.85; Virus x Period: F_(1, 22)_ = 0.04, *P =* 0.84; Training x Virus x Period: F_(3, 66)_ = 0.51, *P =* 0.68. \**P* < 0.05, \*\**P* < 0.01 relative to preCue baseline, Bonferroni correction. ChR2, *N* = 11, 6 males; eYFP, *N* = 13, 6 males.

**Supplemental Figure 4-2.**
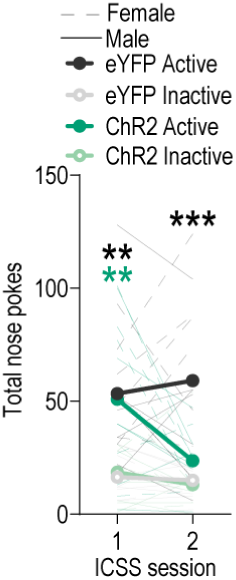
Stimulation of VTA_DA_→BLA projections is not reinforcing. To assess the reinforcing properties of VTA_DA_→BLA activation, rats were given 2 sessions of intracranial self-stimulation (ICSS) in a context different from that of prior conditioning. Nose pokes in the active port triggered 1-s blue light delivery (473 nm; 10 mW; 25 ms pulse width; 20 Hz). Data show total active nose pokes compared to inactive nose pokes across 2, 1-hr ICSS sessions. Activation of VTA_DA_→BLA projections was not reinforcing. Rats expressing ChR2 showed similar levels of active nose pokes as the eYFP control group in the first session and this decreased to the level of the inactive nose pokes in the second session (Session x Virus x Nose poke: F_(1, 22)_ = 5.00, *P =* 0.04; Virus x Nose poke: F_(1, 22)_ = 5.18, *P =* 0.03; Session x Virus: F_(1,22)_ = 5.18, *P =* 0.03; Session x Nose poke: F_(1, 22)_ = 1.24, *P =* 0.28; Session: F_(1, 22)_ = 3.05, *P =* 0.09; Virus: F_(1, 22)_ = 1.94, *P =* 0.18; Nose poke: F_(1, 22)_ = 54.66, *P* < 0.0001). *\*P <* 0.05, *\*\*P <* 0.01 relative to inactive nose pokes, Bonferroni correction. Elevated active v. inactive port nose poking in both the eYFP and ChR2 groups could have resulted from the prior association formed between blue light and reward delivery during compound conditioning. If true, then this could have extinguished by the second session in the ChR2 group, potentially indicating that VTA_DA_→BLA projection activity during either initial learning or online during the ICSS session may contribute to the reward expectation and/or learning processes that contribute to extinction. Alternatively, the nose poking in both groups could reflect salience of the light delivery, which could habituate more quickly in the ChR2 group. ChR2, *N* = 11, 6 males; eYFP, *N* = 13, 6 males.

## Notes

### Competing Interest Statement

The authors have declared no competing interest.

### Summary of Updates

This revision includes new experiments, analyses, figures, and text edits. In particular, we have included 3 new experiments. The first is an additional fiber photometry experiment in which we provide considerable additional information on endogenous profile of dopamine release in the BLA during cue-reward learning using a Pavlovian trace conditioning task. We also provide evidence of BLA dopamine responses to unpredicted rewarding and aversive events. The second is an optical inactivation experiment in which we demonstrate that VTADA->BLA projections are necessary for the formation of the identity-specific stimulus-outcome memories that support sensitivity of cue responses to outcome devaluation. The third is a longer duration inactivation experiment, in which we replicate the previous inhibition result when we inhibit throughout the cue and reward during learning.

